# A novel approach to tagging tubulin reveals MT assembly dynamics of the axoneme in *Trypanosoma brucei*

**DOI:** 10.1101/2025.01.27.634986

**Authors:** Daniel Abbühl, Martina Pružincová, Luděk Štěpánek, Eleonore Bouscasse, Rita Azevedo, Mariette Matondo, Vladimir Varga, Serge Bonnefoy, Philippe Bastin

## Abstract

The protozoan parasite *Trypanosoma brucei* is a mono-flagellated cell during the G1-phase of its cell cycle. In order to duplicate, it assembles a new flagellum alongside the mature one, in which further elongation is prevented. Our group proposed a model where the mature flagellum is locked after construction to full length (Bertiaux et al. 2018) and access of new building blocks for elongation is exclusive to the new flagellum. To test this hypothesis directly, we developed a tool for the inducible expression of tagged tubulin. Alpha-tubulin that was tagged with an intragenic Ty-1-epitope behaved indistinguishable from untagged tubulin. Its incorporation was monitored after inducible expression, to follow the assembly dynamics of microtubules in the cell body, the mitotic spindle and the flagellum. In this study we observed that integration of tubulin occurs at the distal flagellum tip at a linear rate and is indeed restricted to the new flagellum in bi-flagellated cells. This is direct evidence that trypanosomes avoid competition between the two flagella by allowing tubulin incorporation only in the new organelle. However, by tracing flagella over several cell cycles we could also show that mature flagella do not remain locked indefinitely. The restriction is lifted briefly after the bi-flagellated cell has divided and the daughter cell inheriting the old flagellum shows incorporation of newly synthesized building blocks again. It then has to lock again before the cell can assemble a new flagellum. Our findings suggest regular incorporation of tubulin at the tip of previously locked flagella. This evidence was supported with an orthogonal approach, with which we monitored the incorporation of HaloTag-tagged radial spoke protein 4/6. Since flagellum length in trypanosomes is stable, this indicates that the entire axoneme is subject to regular events of transient disassembly followed by assembly at its distal tip.

## Introduction

The flagellum is a membrane wrapped cell protrusion involved in motility, cell morphogenesis and signaling, that is found throughout the eukaryotic lineage. The structural core is called the axoneme, a cylindrical arrangement of nine outer MT doublets that surround a central pair of single MTs. In addition to tubulins, the flagellum contains hundreds of other proteins (Billington et al., 2023; Pazour et al., 2005; Subota et al., 2014). Structure, molecular composition, as well as many processes required for its assembly and maintenance, are highly conserved. As ribosomes are absent from the flagellum, its components need to be actively transported to the distal tip, where they are incorporated (Johnson and Rosenbaum, 1992). This is facilitated by protein complexes also called trains that carry cargo towards (anterograde transport) and back from (retrograde transport) the flagellum tip. The process is referred to as intraflagellar transport (IFT) and is driven by kinesin and dynein motors (Kozminski et al., 1995, 1993).

Despite the common basic architecture, cilia and flagella exhibit variability in structure and length, in relation to their function. Flagellum length can be drastically different between species or even between cell types of the same species. Several models have been proposed to explain how cells control flagellum length. At their heart, most of these models aim to describe how cells manage the availability, transport and incorporation of the flagellum’s most important building component, tubulin. Alpha and beta-tubulin are globular proteins that form heterodimers which are assembled into filaments that get laterally connected to form the hollow tubes called microtubules (Nogales et al., 1998; Tilney et al., 1973). The flagellum extends by the addition of new tubulin at the distal tip (Johnson and Rosenbaum, 1992), while it can also be shortened when tubulin building blocks are removed. The balance-point model describes how the interplay between assembly and disassembly rates governs flagellum length. If the assembly rate exceeds the dis-assembly rate, the flagellum elongates, contrarily shortening occurs when the assembly rate is slower (Marshall and Rosenbaum, 2001). When the two rates are equal, flagellum length is at the balance point and therefore stable. The green alga *Chlamydomonas* regulates the length of their flagella by maintaining a balance between a constant dis-assembly rate and a variable assembly rate that can range between 9 – 24µm/h in elongating flagella (Flavin and Slaughter, 1974) (Rosenbaum 1969). Another model postulates that the length of flagella can be regulated by limiting the pool of building blocks that are available for elongation (limited pool model). Once a cell runs out of tubulin, elongation halts and the length of the flagellum is stable. This mode of assembly is employed for example by *Naegleria fowleri* (Fulton and Walsh, 1980; Goehring and Hyman, 2012). Uncovering the timing and the rate of tubulin addition in the flagellum is therefore crucial to understand how cells regulate flagellum length (Hao et al., 2011; Marshall and Rosenbaum, 2001). Here we investigated the regulation of flagellum length in the protozoan parasite *Trypanosoma brucei*, the causative agent of Human African Trypanosomiasis as well as a substantial burden to livestock in sub-Saharan Africa. They are transmitted between mammalian hosts by the bite of tsetse flies (Glossinidae). Every life cycle stage is flagellated and has a characteristic flagellum length (Rotureau and Van Den Abbeele, 2013). *T. brucei* also poses as a formidable model organism to study flagellum assembly due to its exceptional amenability for reverse genetics, relative ease of culturing, conserved IFT machinery and the fully sequenced genome (Langousis and Hill, 2014). The protein content (∼750 proteins) (Subota et al., 2014) and protein localization inside the flagellum are well understood, especially thanks to the global tagging project Tryptag.org, an invaluable resource, where a majority of proteins encoded in the genome have been tagged and the localization data made publicly available (Billington et al., 2023; Dean et al., 2017; Sunter et al., 2023). Apart from the axoneme the most notable structure inside the trypanosome flagellum is the paracrystalline network called the paraflagellar rod (PFR), which is attached to microtubule doublets 4 and 7 of the axoneme. It is potentially involved in providing structural stability and motility but its precise role remains enigmatic. As construction of the axoneme and the PFR are closely linked, the PFR has been used to describe flagellum assembly dynamics. (Alves et al., 2020; Bastin et al., 1999, 1998; Hughes et al., 2012).

The flagellum is the most distinguished morphological feature of this single celled organism and is attached to the cell body along almost its whole length, apart from the distal tip. In order to divide, the trypanosome needs to assemble a new flagellum (NF) in the same cell in which it maintains an existing one. During NF elongation, the length of the old flagellum (OF) does not change. Once the NF reaches around 80% of the OF’s length, the cell divides and one cell inherits the OF and the other one the NF (Abeywickrema et al., 2019). The NF keeps elongating post cell division to ultimately reach the full length (Robinson et al., 1995). This provides the opportunity to investigate growing and mature flagella in a single cell and raises the question of how these cells manage to grow only one of the two flagella, although they both originate from the same cytoplasm (Bertiaux and Bastin, 2020). One explanation of how these cells deal with assembly of only one flagellum is provided by the Grow-and-Lock model (Bertiaux et al., 2018). It was proposed based on work with the first differentiated life cycle stage found in the insect vector, the procyclic trypanosomes, which will also be used in this study. The Grow-and-Lock model states that once a flagellum has reached its full length, it gets locked. The lock prevents further elongation or shortening, enabling procyclic cells to maintain the OF at a length of ∼20µm while being able to grow the NF independently (Bertiaux et al. 2018).

A first insight into the molecular mechanism governing this process has been revealed with the discovery of the CEP164C protein. It is found at the transition fibers that connect the basal body to the flagellar pocket membrane, exclusively at the base of the OF but not the NF. Upon knockdown of CEP164C, OFs of bi-flagellated cells are abnormally long (up to 30µm) while NFs are made too short. As CEP164C is only found at the base of the OF, this phenotype can be explained by the removal of the locking mechanism leading to unnatural elongation of old flagella. Since new flagella in these cells are also shorter, this likely occurs at the cost of diminished new flagellum elongation. The locking mechanism therefore prevents competition between the two organelles (Atkins et al., 2021).

The Grow-and-Lock model is supported by indirect evidence: When IFT frequency was reduced by RNA interference against IFT kinesins 2A and 2B, the affected cells formed shorter flagella (Bertiaux et al., 2018). However, when de-induction experiments were performed, and IFT frequency was restored to normal, this did not increase the length of the OF. Additionally, in cells where division was inhibited, the NF reached 100% of the OF length but did not exceed it, while in regularly dividing cells, the NF reached about 80% of the OF length before cytokinesis. This suggests that trypanosomes halt flagellum elongation at a certain point to prevent its further growth.

Therefore, we wanted to directly measure if incorporation of new building blocks is indeed prevented in the OF by a locking mechanism and if OFs stay locked forever after they are constructed to full length.

To fully understand how flagella are constructed and maintained during the cell cycle, the ideal situation would be to monitor tubulin incorporation and turnover. So far, the best tool for tracking recently assembled tubulin in *T. brucei* was the Yl1/2 antibody (Kilmartin et al., 1971). It detects the C-terminal tyrosine residue of alpha-tubulin, which is removed (de-tyrosination) shortly after tubulin integration into MTs. Staining with this antibody revealed the signal to be prominent in the distal portion of the growing flagellum and the posterior cell body (Sherwin et al., 1987). However, the window of observing tyrosinated tubulin is restricted to the short time between its polymerization into MTs and de-tyrosination. Furthermore, re-tyrosination could obscure interpretation (van der Laan et al., 2019), although, it seems absent in flagella (Sherwin et al., 1987). For a more long-term picture, it is vital to track tubulin integration flexibly over multiple cell cycles, with each lasting ∼ 9 hours.

Here, we attempted to decipher when during the cell cycle is tubulin incorporated into each flagellum and estimate the rate of incorporation. Towards this goal we attempted to develop an inducible system for expression of tagged tubulin. So far, tagging tubulin in *T. brucei* has proven challenging as tagged tubulin was expressed but not incorporated into MTs (Bastin et al. 1996) or only in a low proportion of cell body MTs but not in the flagellum (Sheriff et al. 2014). Western blot data showed that a large portion of YFP-tubulin was found in the soluble pool and not incorporated into MTs of the cytoskeleton, in contrast to endogenous tubulin (Sheriff et al., 2014).

We have now overcome this problem by tagging *T. brucei* alpha-tubulin with an internal tag inside the acetylation loop situated within the lumen of the microtubule (Sirajuddin et al., 2014; Soppina et al., 2012). After inducible ectopic expression, the incorporation of tagged tubulin could be followed over different periods of time in NFs and OFs during the cell cycle. These experiments demonstrated that tubulin is incorporated at the distal tip of growing flagella at a linear rate and that tubulin integration in bi-flagellated cells indeed follows the locking model. In addition, we managed to track flagella over several cell cycles after their emergence, revealing novel aspects of tubulin incorporation in monoflagellated cells. Moreover, we obtained similar results for another axonemal constituent, the radial spoke protein RSP4/6, indicating that these events likely occur for the other axonemal components as well.

## Results

### Tagged tubulin is incorporated into MTs of the cell body, flagellum and mitotic spindle

Alpha-tubulin was tagged with one or two copies of the Ty-1- epitope (a sequence of 10 amino acids derived from a yeast viral particle) (Bastin et al., 1996) inside the acetylation loop of alpha-tubulin (Schatz et al., 1987; Sirajuddin et al., 2014; Soppina et al., 2012). Fig. S1 illustrates the insertion of the Ty-1-epitope, two amino acids after the acetylated lysine (Fig. S1A, K40 in cyan). In the diploid *T. brucei* genome, alpha and beta-tubulin are found in ∼20 alternating, identical tandem repeats. First, the tagging strategy was tried by *in situ* tagging alpha-tubulin at one of the ∼40 loci (Berriman et al., 2005; Ersfeld et al., 1998; Seebeck et al., 1983) using *in situ* recombination (Kelly et al., 2007). As trypanosomes utilize their gene specific untranslated regions to regulate timing and level of protein expression (Clayton, 2016), we designed a vector that introduces one or two copies of the Ty-1-tag into alpha-tubulin at its endogenous locus with an unaltered alpha-tubulin 3’UTR. This should reduce the risk of artifacts potentially due to untimely expression or unnatural protein levels. The tagged protein will be referred to as Ty-1-tubulin. Supplemental figure 2A depicts immunofluorescence assays (IFA) with an *in-situ* Ty-1-tagged alpha tubulin cell line (one or two copies of the Ty-1-tag) following co-staining with antibodies recognizing the Ty-1 tag (BB2) and alpha-tubulin (TAT- 1).

In both cell lines (one or two Ty-1-epitopes), Ty-1-tubulin is found in the entire cell body, the mitotic spindle and the flagellum, a localization that matches the one revealed by the TAT-1 antibody (Fig. S2A-B). To test whether Ty-1-tubulin was integrated into MTs, we subjected cells to detergent extraction. This treatment retains components integrated into the cell cytoskeleton but not the soluble proteins (Sherwin and Gull, 1989). Upon extraction, Ty-1- tubulin signal remains present in all the aforementioned structures, demonstrating that it is incorporated into MTs (Fig. S2A-B).

To further confirm that the tagged protein is indeed incorporated into MTs, we fractionated cells by detergent treatment and performed western blotting analysis. Protein samples of whole cells, cytoskeleton extracts and detergent-soluble proteins were run on an SDS-PAGE gel and two membranes were stained, one with the BB2 and the other with the TAT-1 antibody (Fig. S2 D-F). A protein was recognized slightly above the 50kDa band marker with the BB2 antibody and was found to be associated to the cytoskeletal fraction, with only a minimal amount found in the soluble fraction. This distribution profile matches the one for alpha-tubulin revealed by the TAT-1 antibody. Surprisingly, despite being only 1kDa heavier, the tagged protein could be discriminated from the endogenous alpha-tubulin (expected size of alpha tubulin: ∼49.8kDa) as an extra band detected by TAT-1 (Fig. S2 D-F). These results prove that cells were able to readily incorporate alpha-tubulin that contains a Ty-1-epitope inside the acetylation loop into MTs of the cell body, flagellum and the spindle, making it a suitable tag in 1. *T. brucei*.

However, this system is constitutive and does not allow modulation of timing of gene expression. We therefore developed a cell line that allowed inducible ectopic expression of Ty- 1-tubulin (Fig. S1D), which is induced by the addition of tetracycline (Wirtz and Clayton, 1995). Figures 1 and 2 depict a set of validation experiments performed with the derived cell line. Whole cells were grown in the absence (Fig. 1A, non-induced, upper panel) or in the presence of 1µg/ml tetracycline for 16h (Fig. 1A, induced, lower panel) and treated for IFA with the antibody that recognizes the Ty-1-epitope (BB2) as well as DAPI. In the absence of tetracycline, there is no signal in almost all cells (Fig.1A, top right). By contrast, in induced cells tagged tubulin is observed in the entire cell body (Fig. 1B) as well as in the mitotic spindle (Fig. 1B, red arrow) and the flagellum (Fig. 1B, white arrows).

**Fig. 1:**
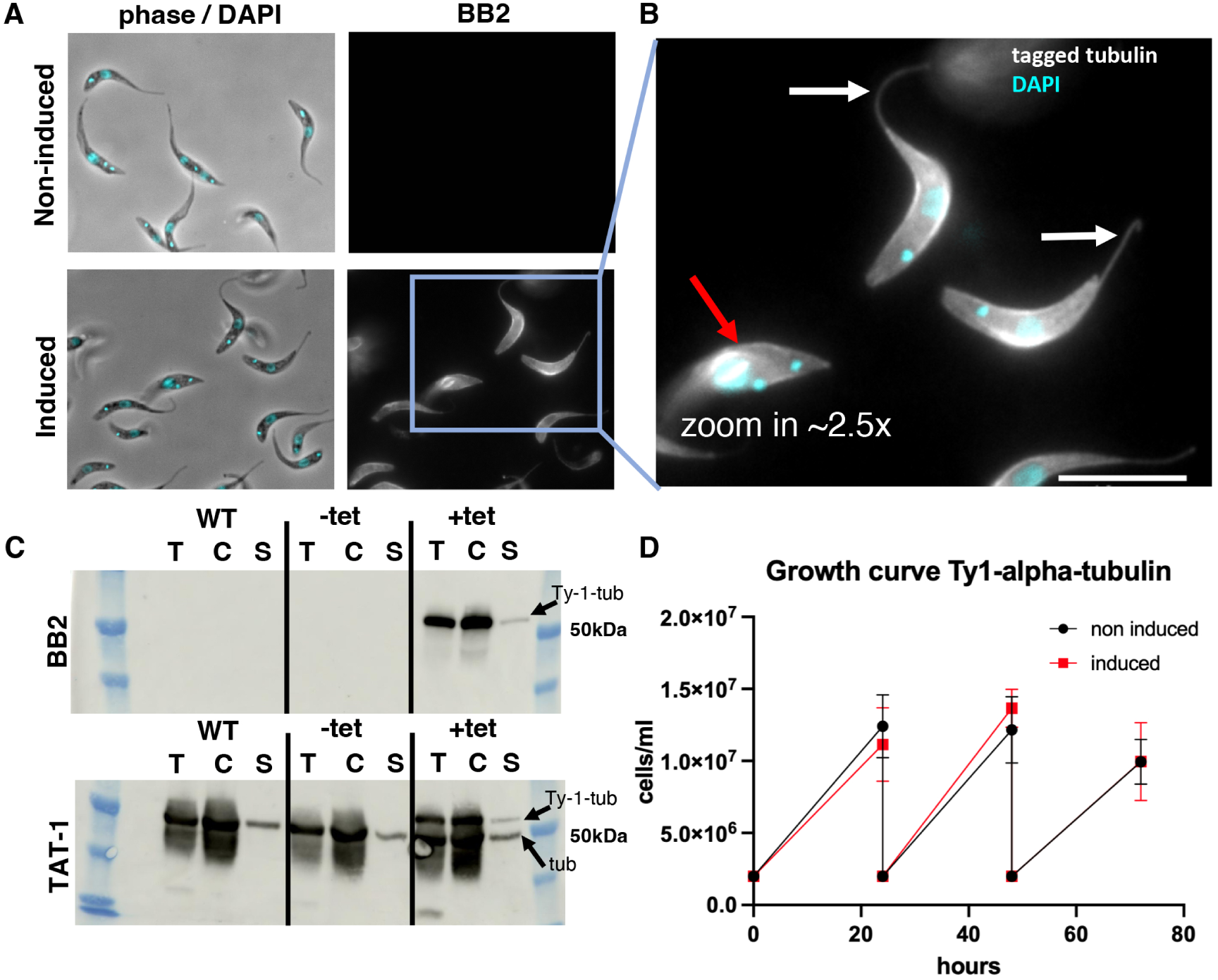
Tagged tubulin is incorporated into MTs. A: IFA with whole cells of cell line capable of an inducible expression of tagged tubulin. Cells were fixed with methanol and stained with the BB2 antibody (Ty-1-tubulin, white) and DAPI (cyan). Top: non-induced, Bottom: Induced with tetracycline for 16 hours. B: Zoom in of induced cells highlighting the presence of tagged tubulin into cell body, mitotic spindle (red arrow) and the flagellum (white arrows). C: Western blots with protein samples of different fractions from the wild-type cell line, the non-induced (-tet) and induced (+tet) tagged tubulin cell line. Top: membrane was stained with BB2 (tagged tubulin). Bottom: membrane was stained with TAT-1 (alpha-tubulin). T: whole cell isolates, C: detergent extracted cytoskeletons, S: soluble proteins. D: Growth curve of non-induced (black circles) and induced (red squares) tagged tubulin cell line. Scale bar = 10µm.

**Fig. 2:**
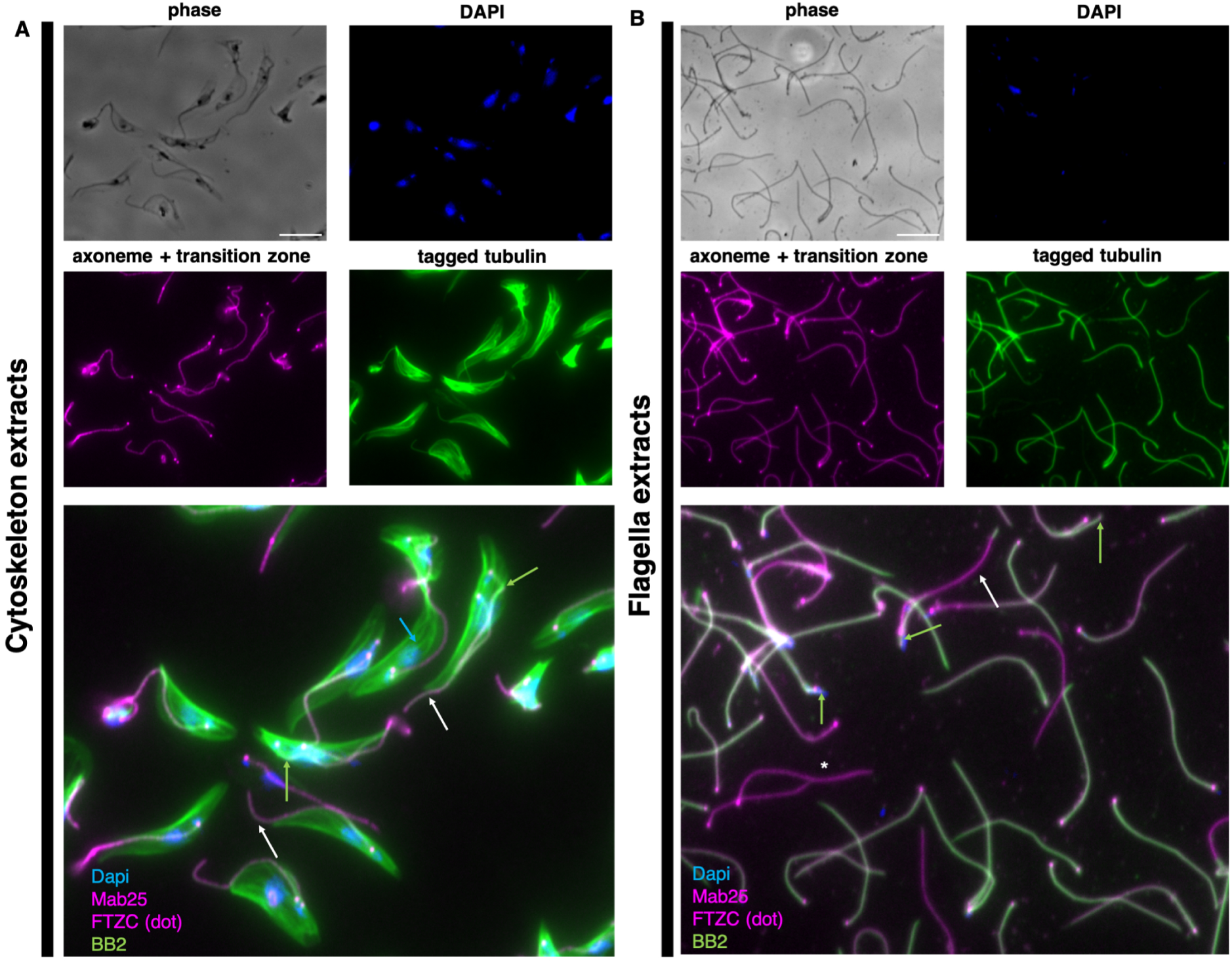
**Ty-1-tubulin is incorporated into MTs of the cell body and the flagellum** IFA with cells 24h after the addition of tetracycline. Samples were incubated with antibodies targeting the axonemal component TbSAXO (Mab25, magenta), the transition zone protein FTZC (anti-FTZC, magenta dot), the antibody that recognises the Ty-1-epitope inside tagged tubulin (BB2, green), and DAPI. A: Whole cells were treated with 0.5% NP40 to obtain detergent extracted cytoskeletons. Ty-1-tubulin is present inside the subpellicular MT corset as well as the flagellum. B: Cytoskeleton extracts were further treated with 1M NaCl to depolymerise the corset MTs to obtain 1M NaCl extracted flagella. Ty-1-tubulin is still present inside flagella and therefore integrated into the MTs of the axoneme. Green arrow: NF, white arrow: OF, blue arrow: cell corset, white asterisk: tetracycline non-responders. Scale bar = 10µm.

To evaluate the proportion of tagged tubulin that remains associated to the cytoskeleton, cell fractionation was performed and analysed by western blotting. In Fig. 1C, two membranes are shown with samples from the wild type cell line (WT), from the non-induced control (-tet), and from cells that were grown for 16 hours with tetracycline (+tet). When probed with the tag specific BB2-antibody, no signal is detected for WT (Fig. 1C, upper, WT) or non-induced samples (Fig. 1C, upper, -tet). In the induced samples (Fig. 1C, upper, +tet), intense bands are visible for the whole cell as well as the cytoskeleton fraction and a band of relatively lower intensity is visible for the soluble pool fraction, as expected (Sherwin et al., 1987). All bands migrated slightly above the 50kDa band of the marker. In the blot stained with the alpha-tubulin antibody, (Fig. 1C, TAT-1) a band that migrated at the size of the 50kDa-marker is visible for all three fractions of all conditions, with the band in the soluble protein fraction being fainter, as expected. Under induced conditions (Fig. 1C, lower, +tet) an additional set of bands is visible slightly above the 50kDa marker, which corresponds to Ty-1-tubulin. Importantly, their distribution profile is comparable with that of untagged tubulin, confirming that the majority of Ty-1-tubulin had integrated into MTs and behaves as endogenous tubulin (Fig. 1C). Finally, this ectopic expression of Ty-1-tubulin does not affect cell growth even when cells were induced for several days (Fig. 1D). We did not observe any aberrancies of the cell cycle when induced and non-induced cells were quantified.

To complement the data obtained from western blots, Fig. 2 depicts two IFA experiments with cells that were harvested 24h after the addition of tetracycline. Whole cells were treated with 0.5% Nonidet P-40 to obtain detergent extracted cytoskeletons and processed for IFA with the BB2 antibody in combination with the mAb25 antibody (anti-TbSAXO protein), which is a marker of the axoneme (Dacheux et al., 2012), and the anti-FTZC antibody (Flagellum Transition Zone Component), a marker of the flagellum base (Bringaud et al., 2000). The tagged tubulin remained associated to the cytoskeleton (Fig. 2A). To better visualize the flagellar skeleton, cytoskeletal extracts were further treated with 1M NaCl, which depolymerizes the MTs of the corset while leaving the ones of the axoneme intact (Robinson et al. 1990). Ty-1-tubulin is indeed associated to the flagellum throughout its length, further confirming its tight incorporation (Fig. 2B). As a fraction of cells did not respond to tetracycline some flagella remained unstained by Ty-1-tubulin (Fig. 2B, asterisk). We also noted some heterogeneity in signal intensity between cells. As we did not observe this when Ty-1-tubulin was constitutively expressed, this is most likely the consequence of increased Ty-1-tubulin levels with time elapsed after induction.

Lastly, we investigated tagged alpha-tubulin for the presence of classic post-translational modifications, especially acetylation since the Ty-1 tag is positioned very close to the acetylated lysine K40 (Fig. S3A). To assess the acetylation status of Ty-1-tubulin, protein isolates of an induced culture (7 days) were compared to a non-induced control using western blotting. Membranes were stained with TAT-1 and Yl1/2 (that detects tyrosinated alpha-tubulin) revealing the presence of endogenous and tagged tubulin, the latter one migrating a bit more slowly (arrows in Fig. S3A). Then, two distinct antibodies that recognize acetylated alpha tubulin were used (C3B9 and 6-11B-1 (Piperno and Fuller, 1985; Woods et al., 1989)), but they detected only the endogenous tubulin (Fig. S3A). According to literature, however, virtually all incorporated tubulin is acetylated (Schneider et al., 1997). As Ty-1-tubulin behaves in other respects indistinguishable from untagged tubulin, the absence of this modification was unexpected. We reasoned that the insertion of the tag just two amino acids after lysine 40 could destroy the binding epitopes of C3B9 and 6-11-B-1. Therefore, the Ty-1-tubulin cell line was induced for 7 days, cells were harvested and their flagellar skeletons isolated by detergent treatment followed by exposure to 1M NaCl as above (Robinson and Gull, 1991). These isolates were then subjected to analysis by mass spectrometry after digestion with three different proteases. Thanks to the close presence of the tag (Fig. S3B), peptides derived from the Ty-1 tagged protein can be discriminated from those originating from endogenous tubulin. These experiments showed that acetylation was detected on lysine 40 of peptides derived from both endogenous and tagged tubulin by measuring the presence of acetylated peptides after AspN cleavage (four replicates shown in Fig. S3C). This confirmed that Ty-1-tubulin is acetylated and that detection by antibodies likely failed due to the alteration of their binding epitope.

Lastly, the timing of Ty-1-tubulin expression upon tetracycline addition was evaluated by western blotting, revealing that it could be observed as early as one-hour post induction, with a rapid increase over time (Fig. S4A-B). The bands corresponding to the soluble pool of Ty-1- tubulin were relatively faint, indicating that the majority was found in the cytoskeleton fraction and therefore incorporated into MTs, even at early time points of induction. As a full cell cycle lasts around 8-9h, this is sufficient to measure assembly dynamics of the axoneme during NF construction. We conclude that we have succeeded to tag alpha-tubulin in a way it gets properly incorporated into all the MTs of trypanosomes. Combined with the rapid expression provided by our inducible system, this offers the possibility to directly monitor tubulin incorporation into trypanosome flagella for the first time.

### Tagged tubulin is distally integrated into the axoneme at a linear rate

The assembly rate of the *T. brucei* axoneme has so far been estimated indirectly, based on tubulin detyrosination (Sherwin and Gull, 1989), cell cycle duration (Woodward and Gull, 1990) or the measurement of an extra-axonemal component (Bastin et al., 1999). To analyse the kinetics of tubulin integration into the axoneme, we performed time courses by inducing expression of Ty-tubulin and taking samples at various points of induction.

At first, Ty-1-tubulin expression was induced for short periods of 1-4 hours and the incorporation was monitored at the level of the entire cell. From previous studies, it is known that newly synthesized tubulin incorporates into subpellicular microtubules at the posterior cell body end (Sherwin et al., 1987)(Sheriff et al., 2014) and the same pattern was observed upon short induction times of Ty-1-tubulin (Fig. S5A). In addition, newly synthesized Ty-1-tubulin is clearly present in the growing flagellum (Fig. S5A, arrows) and in the mitotic spindle (Fig. S5A, stars). To visualize flagella more clearly, subpellicular MT were depolymerised with 1M NaCl treatment (Fig. S5B). In these flagella isolates, Ty-1-tubulin is found integrated exclusively in the distal portion of the new flagellum. This is a direct proof that the addition of the new tubulin occurs at the distal tip of a growing flagellum in *T.brucei*.

As Ty-1-tubulin localization after tetracycline addition follows the expected patterns, we were able to determine the kinetics of Ty-1-tubulin integration into the axoneme of the new flagellum during its assembly. This was done by measuring the length of flagellar segments that had incorporated Ty-1-tubulin after different periods of induction. As trypanosomes divide asynchronously and all cell cycle stages are present in a culture (Sherwin and Gull, 1989; Woodward and Gull, 1990), we focused our measurements on cells with a new flagellum that had integrated Ty-1-tubulin in a distal segment. Time courses were performed, where tagged tubulin expression was induced between 2-4 hours and samples were then taken for analysis every 30 minutes. As tubulin signal in flagella is sometimes hard to distinguish from signal of the MTs in the cell body, we decided to analyze the assembly rate with 1M NaCl-extracted flagella. Conveniently, procyclic trypanosomes possess a structure called the flagella connector (FC), that links the tip of the NF axoneme to the side of the OF axoneme (Briggs et al., 2004). The FC is resistant to detergent and 1M NaCl treatments upon which the connection of the NF to the OF is maintained (Moreira-Leite et al., 2001)(Fig. 3A). To highlight the proximal end of the axoneme, the transition zone marker FTZC was used. It can be observed as a dot that marks the very proximal portion of the flagellum. The new flagellum is found in a posterior position, is shorter and is connected at its tip to the OF axoneme.

**Fig. 3:**
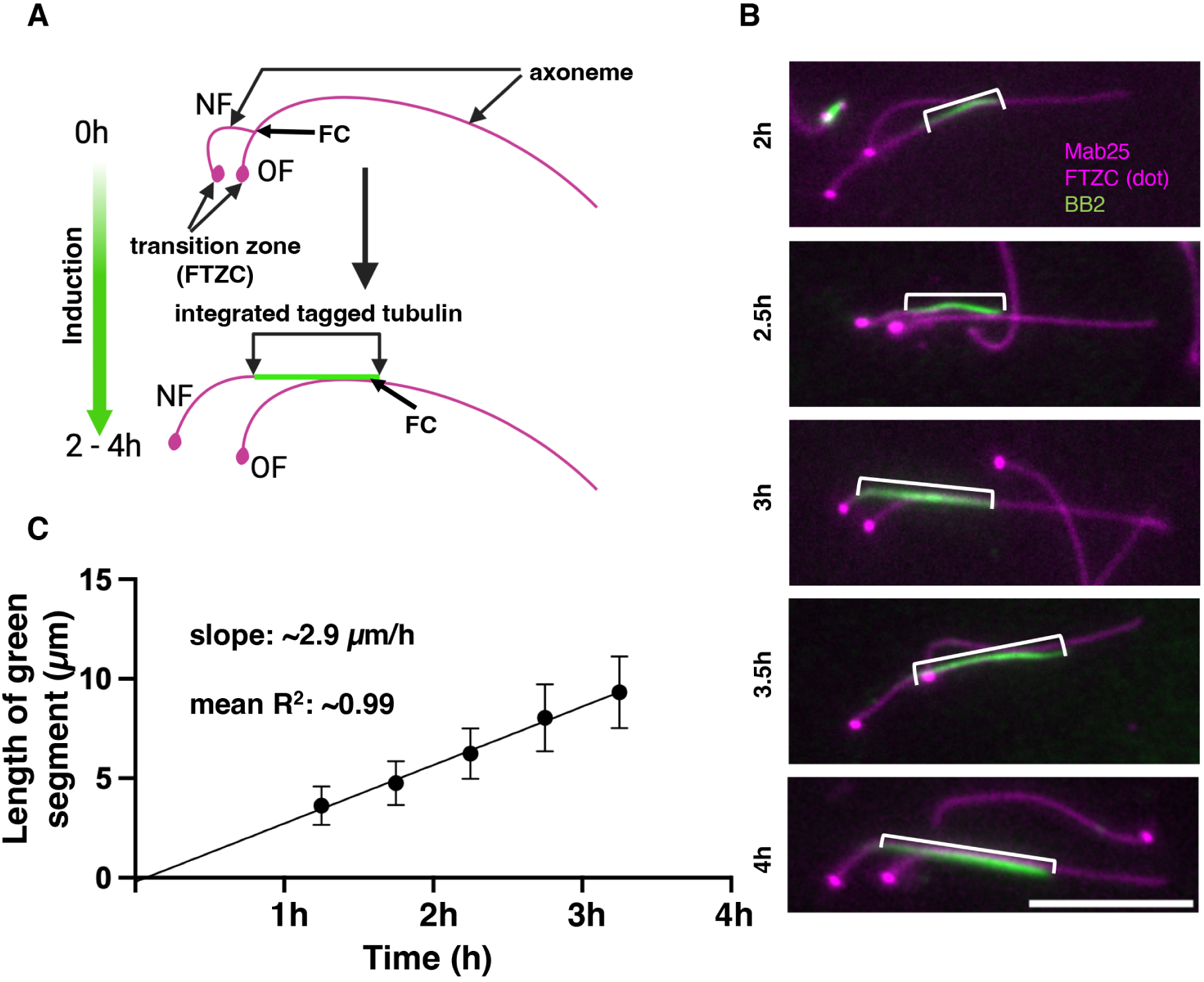
**Tagged tubulin is incorporated at a linear rate into the new flagellum.** A: Cartoon depicting isolated flagella before and after induction of Ty-tubulin expression (2-4h). This cell has started NF assembly before induction. The dot in magenta marks the transition zone (proximal portion of the flagellum) and the line in magenta is the axoneme. Top: A cell before induction with a NF during assembly and with an OF. Bottom: The same cell after induction. If integration takes place at the distal end as expected, the tagged tubulin synthesized after the addition of tetracycline should integrate as highlighted in green. B: Example images of flagella isolated from five timepoints post induction. Flagella were extracted with 1M NaCl concentration and incubated with antibodies targeting the axonemal component TbSaxo (Mab25, magenta), the transition zone protein FTZC (anti-FTZC, magenta dot), the antibody that recognises the Ty-1-epitope inside tagged tubulin (BB2, green). White spacers highlight the portion of the NF in which tagged tubulin has integrated. C: The length of the green portion highlighted in Fig. 2B was measured. Three biological replicates (70 - 100 flagella each) were pooled and plotted. Timepoints were corrected by subtracting 45 minutes from the induction time, which is an estimated response time for tetracycline addition to protein expression (Wirtz and Clayton 1995). Regression line shows linear growth with a 99% confidence interval. Scale bar = 10µm.

A cartoon of flagella of a cell that had started assembly of its new flagellum before addition of tetracycline is depicted in Fig. 3A, 0h. Once Ty-1-tubulin is produced, it should be added at the distal tip of the growing NF (Fig. 3A, 2-4h, Fig. S5B). Individual examples are shown for each timepoint in Fig. 3B, where the Ty-1-tubulin containing segment that was measured is indicated with the white bracket. A gradient of increasing intensity of Ty-1-tubulin signal towards the distal tip was noted. This was expected as Ty-1-tubulin becomes relatively more abundant during the course of induction (Fig. S4, Fig. 3B).

The length of the Ty-1-tubulin containing segment of these NFs was measured and the data of three separate experiments (70 - 100 flagella per timepoint and experiment, ∼1500 total) was pooled (Fig. 3C). A linear regression was performed for the five timepoints (2-4 hours, 30 minute intervals). This revealed that tagged tubulin was added to the distal tip of NF at a linear rate of ∼2.9µm/h. The linear regression line in Fig. 3C was corrected by subtracting 45 minutes from each timepoint due to the previously reported delay between addition of tetracycline and the expression of the respective protein (Wirtz and Clayton, 1995). We conclude that the flagellum in procyclic trypanosomes is constructed at a constant rate and that our expression system is a suitable tool to dynamically track flagellum assembly during the cell cycle.

### Old flagella of bi-flagellated cells do not incorporate tagged tubulin at early timepoints

We next investigated in which flagella tagged tubulin had incorporated during 2-4 hours of induction. According to the Grow-and-Lock model, OF are not expected to integrate tagged tubulin as the lock supposedly prevents further extension or shortening (Bertiaux et al., 2018; Bertiaux and Bastin, 2020). However, one could not rule out a perfect balance between assembly and disassembly as it is observed in *Chlamydomonas* (Marshall and Rosenbaum, 2001), which would result in no net change in flagellum length. We therefore examined bi- flagellated cells at different timepoints following tetracycline addition for the presence of integrated Ty-1-tubulin in their OF. Cells were induced for 2-4h and harvested every 30 minutes. Flagella were extracted with 1M NaCl and analysed as above.

Cartoons in Fig. 4A and B illustrate two possible cases according to the stage of the cell cycle: in the first case (Fig. 4A), NF assembly had started after the addition of tetracycline and the NF therefore incorporated tagged tubulin along its entire length. In the second case (Fig. 4B) the NF had emerged before inducible expression and Ty-1-tubulin is therefore absent in the proximal portion of the NF. In both cases, two scenarios are possible for the OF, either Ty-1- tubulin is found integrated at the distal tip of the OF as well (scenario 1) or not (scenario 2).

**Fig. 4:**
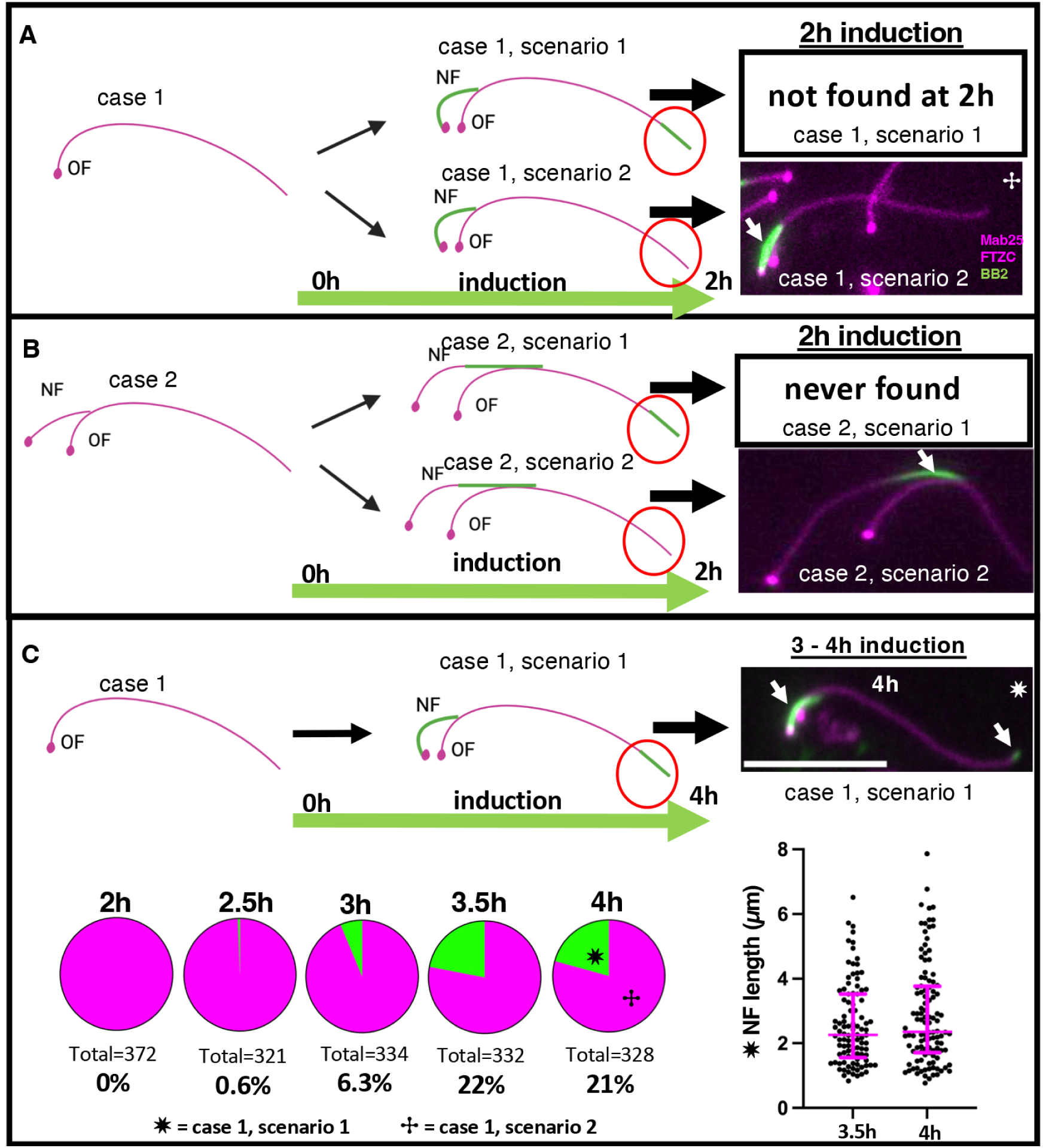
**At early timepoints of induction tagged tubulin is incorporated only in NF of bi- flagellated cells.** A: The cartoons depict two possible scenarios of tubulin assembly into flagella before (left) and after induction (middle) and corresponding flagella that were observed by IFA (right). Assembly of NF started after induction and NF therefore incorporated tagged tubulin along the entire length (case 1). The upper cell is proposed to have incorporated tagged tubulin in the NF and at the tip of the OF (case 1 scenario 1). At 2 hours post induction no such cells were found. Lower: The cell incorporated tagged tubulin only in the NF but not the OF (case 1 scenario 2). At 2 hours post induction these cells were frequently observed. B: Cartoon depicts two possible scenarios of tubulin assembly into flagella in a cell, which started NF assembly before induction (case 2). Situation before (left) and after induction (middle) and corresponding flagella that were observed by IFA (right). As assembly of NF started before induction, the proximal portion has incorporated only untagged tubulin (case 1). The upper cell incorporated tagged tubulin in the distal portion in both OF and NF (case 2 scenario 1). No such cells were found at any timepoint after induction. The lower cell has incorporated tagged tubulin in the distal part of the NF, but not the OF (case 2 scenario 2). These cells were frequently observed. C: At later timepoints of induction, cells were observed that had integrated tagged tubulin along the entire length of the NF (white arrow) but also at the very distal tip of the OF (white arrow, case 1 scenario 1). The frequency of these cells is shown in pie charts (green). Bottom right: NF length of cells with Ty-1-tubulin integrated along the entire NF. ✷ = case 1 scenario 1, ✢ = case 1 scenario 2, Scale bar = 10µm.

When we focused on relatively short NFs (2-5µm) after 2 hours of induction, Ty-1-tubulin had integrated along the entire length as expected. At this timepoint, integration of Ty-1-tubulin in OF from bi-flagellated cells was not observed in any of the >300 OFs that were analyzed, across three separate experiments.

In the second case of cells illustrated in Fig. 4B (cartoons), assembly of the NF had started before the addition of tetracycline. One would expect that Ty-1-tubulin is incorporated at the distal portion while the proximal portion in the NF should contain only untagged tubulin. In comparison to cells from Fig. 4A, these NFs were longer and Ty-1-tubulin was indeed found integrated in the distal segment (Fig. 4B, case 1 scenario 2). However, simultaneous distal incorporation of Ty-1-tubulin in both NF and OF was never observed (N>1500) (Fig. 4B, case 2 scenario 1). Hence, from these observations we concluded that during assembly of NFs, before cytokinesis, the OFs do not extend.

When we induced cultures over longer periods (3 – 4h), a different profile emerged for case 1 cells: we observed bi-flagellated cells where tagged tubulin had integrated along the entire length of the NF (case 1), but also at the distal tip of the OF (scenario 2) (Fig. 4C arrows), which was not observed after 2 hours. The portion of the OF containing tagged tubulin was typically discrete and short. Over time the number of these cells progressively increased to ∼22% at 3.5 - 4h post induction (Fig. 4C). Presence of Ty-1-tubulin in OF was exclusively found in cells where the NF was relatively short (∼2.5 – 2.8µm average length, Fig. 4C, bottom right), and fully labelled with Ty-1-tubulin.

By contrast, when Ty-1-tubulin had integrated only in the distal portion of the NF (case 1) it was never found in the OF (Fig. 4B, case 1 scenario 1). This held true for any timepoint between 2 – 4h of induction.

This meant that integration of Ty-1-tubulin in OF and then NF occurred in succession and not simultaneously, marking the first direct demonstration for the locking mechanism that prevents integration of new tubulin into the OF. The fact that Ty-1-tubulin had only integrated in the distal portion of the OF in cells that carried a short fully labelled NF at later time points suggests that this OF portion might correspond to the last elongation step of a flagellum in the NF daughter. Indeed, assembly of the flagellum in the daughter cell is only completed after cell division (Abeywickrema et al., 2019; Bertiaux et al., 2018; Bertiaux and Bastin, 2020; Farr and Gull, 2009). We investigated this possibility in the following experiments.

### Old flagella frequently integrated tagged tubulin at the distal tip after cell division

As integration of recently synthesized tubulin in OFs of bi-flagellated cells was only observed from 3-4h post induction, we decided to investigate the exact origin of this signal and what it might represent. The cartoon in Fig. 5A illustrates the progression of the cells in scenario 2 (Fig. 4A) over the duration of one cell cycle (∼9 hours) after the addition of tetracycline. The NF in the illustrated example (Fig. 5A) has emerged before induction. It then progressively incorporates Ty-1-tubulin (in green) at the distal tip (4h). At the expected 5h mark, the cell divides, resulting in two mono-flagellated daughter cells in G1-phase. The daughter cell inheriting the NF has integrated Ty-1-tubulin in a distal segment of its flagellum; we refer to these flagella as first generation-OFs (Fig. 5A, ^1st-gen^OF). The OF daughter (Fig. 5A, OF) does not incorporate Ty-1-tubulin, similar to what was shown in Fig. 4B, in agreement with the Grow-and-Lock model (Bertiaux et al., 2018).

**Fig. 5:**
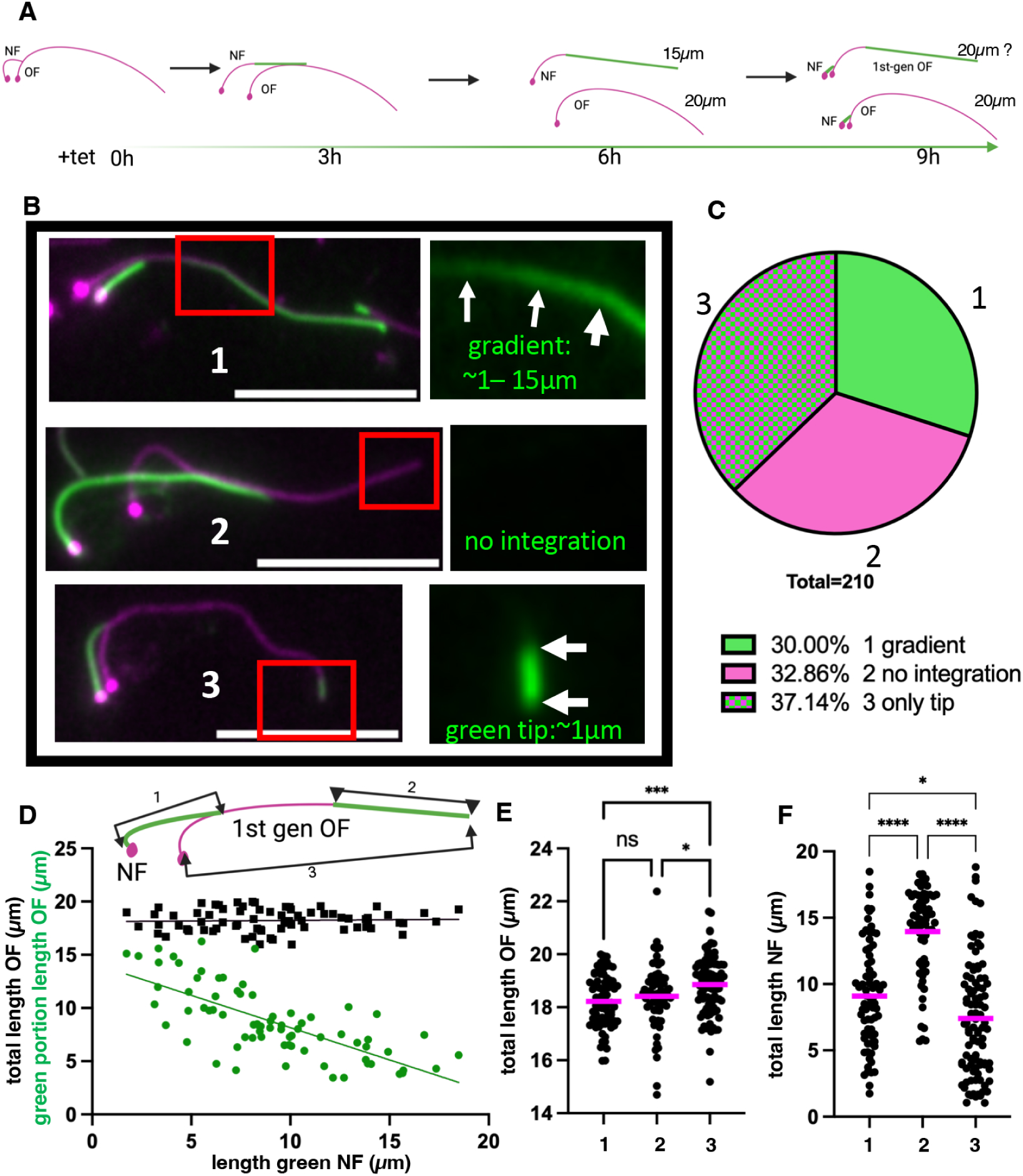
**Tagged tubulin is frequently incorporated into the OF tip.** A: Cartoon shows the progression of a cell that had started the assembly of a NF before induction over the length of one cell cycle (9-hour induction). While the NF is assembled, tagged tubulin is integrated in the distal portion. Over time the cell will divide and one cell will inherit the NF (top, 6 and 9h) and one cell will inherit the OF (bottom, 6h and 9h). Both daughter cells will then start to assemble a NF again. These NF will be fully green. The OF of the NF daughter (1^st^ gen OF, 9h, upper cell) can be distinguished from the OF of the OF daughter as it has a green gradient of tagged tubulin with increasing intensity towards the distal tip. **B:** Examples of cells with a green NF grouped by the different types of OF. 1: A new ^1st-gen^OF OF (Cartoon A 9h top,). 2: An old flagellum of an OF daughter (Cartoon A 9h bottom,). 3: OF that had integrated tagged tubulin exclusively at the very distal tip (average length ∼1µm). White arrows = Ty-1-tubulin in OF. **C:** Frequency of bi-flagellated cells grouped by the different types of **D:** Black squares: the total length of type 1 cell OF (Y-Axis, segment 3) plotted versus the length of the corresponding NF (X-Axis, segment 1) in the same cell. Green dots: the length of the green portion in the OF (Y-axis, segment 2) plotted versus the length of the NF (X-Axis, segment 1). **E:** Comparison of the length of OF grouped by the different OF types described in B. **F:** Comparison of the length of NF grouped by the different types of the OF. Scale bar = 10µm.

After 9h in the presence of tetracycline, both daughters will have started the assembly of a NF that should incorporate Ty-1-tubulin along its entire length. Hence, Fig. 5A illustrates how ^1st- gen^OF can be used to dissect the timing of Ty-1-tubulin integration as the time of their emergence can be precisely determined in respect to the start of the induction. To establish if cells indeed follow this model, we induced the Ty-1-tubulin cell line for 9 hours (∼1 cell cycle). Cells were then harvested and 1M NaCl flagella were stained with BB2, Mab25 and α-FTCZ as above. Cells with a fully labelled NF were then analyzed for Ty-1-tubulin integration in the corresponding OF. We observed that the staining intensity profile of the OF could be separated into three distinct categories (Fig. 5B). The first type corresponds to the ^1st-gen^OF (see Fig. 5A, 6 and 9h) as evident by the presence of the Ty-1-tubulin only in the distal segment. The Ty-1- tubulin containing segment has a clear gradient of increasing intensity towards the tip (Fig. 5B, type 1). As described in Fig. 3, we attribute this gradient to the increasing relative amount of Ty-1-tubulin over the course of flagellar elongation. The second type corresponds to OF in which Ty-1-tubulin is entirely absent from the OF (Fig. 5B, type 2). According to the Grow-and-Lock model, this should be the case for all flagella that were OF at the time of induction. Surprisingly, a third category of OFs was found in which Ty-1-tubulin had integrated only in the very distal tip. This Ty-1-tubulin containing segment has a mean length of ∼1µm (Fig. 5B, type 3) and the transition between it and the preceding segment made of untagged tubulin was abrupt (Fig. 5B type 3, white arrows) rather than in a form of a gradient, in contrast to type 1 OF (see white arrows in magnified panels at the right of Fig. 5B). According to the Grow-and-Lock model and to the linear growth rate of the flagellum, an integration of 1µm should correspond to the last ∼5% of the elongation of a NF to its maximum length (∼20µm). If this 1µm segment indeed corresponded to this final step of linear growth, a low number of type 3 flagella would be expected (∼7.5%). An estimate of the abundance of each OF type according to the Grow-and-Lock model is shown in Fig. S6.

The relative abundance of these 3 types was quantified (Fig. 5C). Type 1 OF constituted ∼30%, type 2 OF ∼33%, and type 3 OF ∼37% of bi-flagellated cells. Such a high number of type 3 flagella is surprising and is indicative of a distinct event that occurs more frequently, than the construction of the last 1µm segment via linear growth.

In shorter induction time point experiments (Fig. 4) simultaneous integration of Ty-1-tubulin in NF and OF was not observed, which suggested that cells had completed OF elongation before the assembly of a NF and that no further tubulin incorporation into OF was taking place. We therefore expected a relationship between the length of the labeled segment in the OF and the length of its corresponding NF. For each cell the length of the Ty-1-tubulin containing segment of the OF (Fig. 5D, segment 2) was plotted against the length of the NF (Fig. 5D segment 1). A clear negative correlation can be observed (Fig. 5D, green dots, segment 1 vs. 2). Additionally, the average cumulative length of the fully labeled NF and the labeled segment in the OF was ∼18µm. This means that an average cell spent approximately 6 hours assembling a NF at a linear assembly rate of 3µm/h (6h x ∼3µm/hour = ∼18µm), including the elongation that occurs after cytokinesis. The total length of the OF (Fig. 5D, segment 3) however did not show a correlation with that of the NF (Fig. 5D, segment 1).

The question when new flagella reached their final full length after cell division remains open (Farr and Gull, 2007). To assess if ^1st-gen^OF (type 1) have reached full length before the bi- flagellate stage, their length was compared to the length of the two other types of OF (Fig. 5E). On average the length of type 1 OF (by definition younger since they were assembled during the induction period) is slightly but significantly (ordinary one-way Anova) shorter when compared to OF that had integrated Ty-1-tubulin at the distal tip (type 3). This length difference of ∼0.8µm was similar to the average length of the green tip (∼1µm, Fig. 5B, type 3, right panel). As assembly of NF is linear (at least during the bi-flagellated stage, Fig. 3C), the length of the NF is a reliable indicator of how advanced an individual trypanosome is in its cell cycle (Robinson et al., 1995). Fig. 5F shows the average length of the NF in cells sorted by the type of their corresponding OF. For youngest OF (type 1), the length of the NF follows a normal distribution per expectation, as the ^1st-gen^OF correspond to cells with an assembling NF of any length at the time of induction. NF length at any given point in time in a exponentially growing culture follows Gaussian distribution (Woodward and Gull, 1990), in agreement with the linear growth rate measured here. In OF where no tagged tubulin integrated (type 2), the corresponding NF was on average significantly longer (ordinary one-way ANOVA), which indicated that integration might be linked to an event at the very beginning of G1 after cytokinesis. NF length of type 3 flagella was broadly distributed but nonetheless tending to be shorter compared to NFs in cells type 1 and type 2 OFs.

Together, these results suggest that integration of tagged tubulin in the distal OF tip occurs frequently but is restricted to a certain window in the cell cycle. Judging by the distribution of NF lengths in type 2 (absence of incorporation) and type 3 (presence), this might occur early on in G1-phase.

### Integration of Ty-1-tubulin during the G1-phase

In the previously described experiments, we observed that ^1st-gen^OF (type 1, Fig. 5) are on average slightly shorter compared to the other two types (Fig. 5E) and that ∼37% of OF had integrated Ty-1-tubulin only at the very distal tip (Fig. 5C, type 3). It is conceivable that this integration corresponds to a distinct elongation step that ^1st-gen^OF (type 1) would have to undergo in order to reach maximum length. Another possibility is that the integration at the tip corresponds to a more frequently occurring event, that happens in all flagella, presumably when cells are monoflagellated. This would entail that the locking mechanism is briefly removed to allow integration during G1-phase. It would then have to be re-applied before the cell initiates the assembly of a NF. This is coherent with the localization pattern of CEP164C, which is absent in 60% of monoflagellated cells (Atkins et al., 2021).

In the following experiments, we tracked flagella for longer periods to investigate if they were subject to integration of tubulin over multiple cell cycles following their emergence. A pulse-chase experiment was designed in which cells were initially pulsed with a 24-hour induction with tetracycline corresponding to ∼2-3 cell cycles and then subjected to a 48h de-induction. These pulse-chase experiments are presented in Fig. 6 (flagellar skeletons) and Fig. S8 (detergent extracted cytoskeletons).

**Fig. 6:**
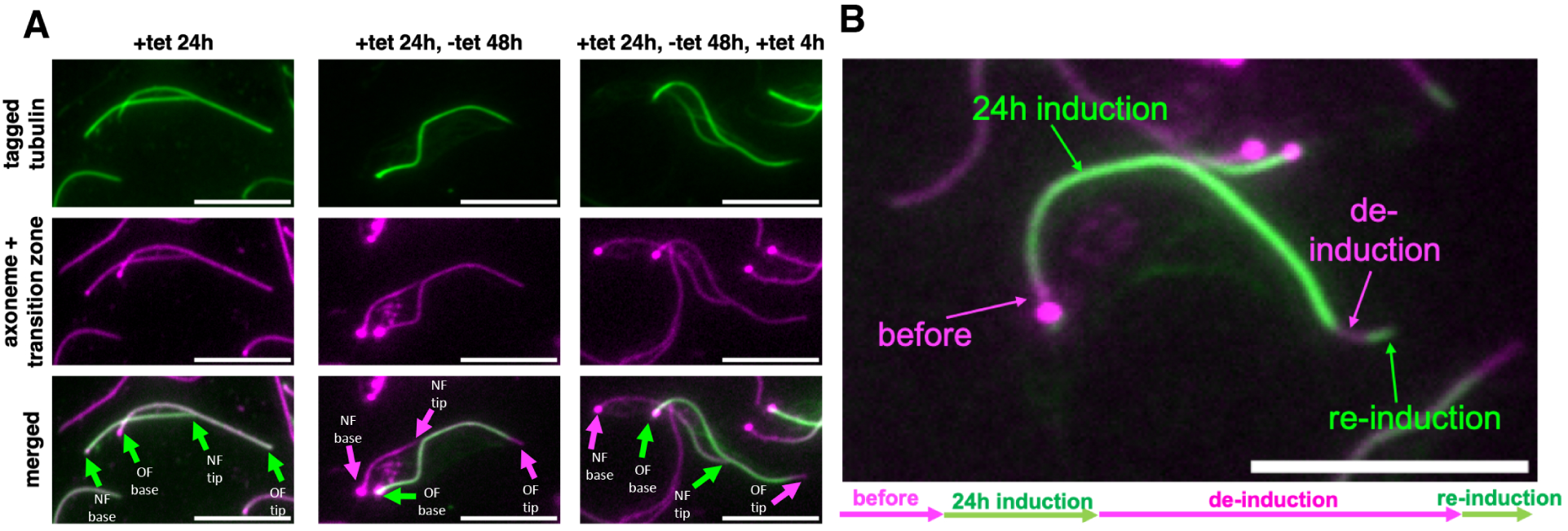
**Tagged tubulin is frequently incorporated into the OF tip in cells that are in G1-phase.** A: Flagella extracts of cells that were subject to a pulse-chase experiment. Cells were first induced with the addition of tetracycline to the medium for 24h (left panel, +tet 24h). Tetracycline was then washed out and the cells were harvested after being grown in absence of tetracycline for 48h (middle panel, +tet 24h, -tet 48h). After that the cells were re-induced with tetracycline for 4h and harvested for IFA (right panel, +tet 24h, -tet 48h, +tet 4h). Samples were stained with BB2, Mab25 and anti-FTZC. Green arrows highlight the presence of Ty-1- tubulin at the flagellum base and tip. Magenta arrows highlight the absence of Ty-1-tubulin. B: An example of flagella from mono-flagellated cells (G1-phase). Arrows indicate the times during the pulse chase experiments in which the corresponding part was most likely assembled. Scale bar = 10µm.

After 24h in tetracycline cells were harvested and whole cells, detergent extracted cytoskeletons and flagella were isolated and subjected to IFA. The samples were stained with BB2, Mab25, α-FTZC antibodies and DAPI as above. We observed that ∼95% of cells had integrated Ty-1-tubulin into MTs of the cell body or flagella and were therefore responding to tetracycline (Fig. S7A).

In preparations of extracted flagella, we scored OFs of bi-flagellated cells and the single flagellum of monoflagellated cells. We observed that a majority (∼63%) of them had integrated tagged tubulin along the entire length, 20% had incorporated Ty-1-tubulin at the tip and ∼11% showed a gradient (Fig. S7B). 7% of flagella did not incorporate any Ty-tubulin. This is just marginally higher than the 5% whole cells not responsive to tetracycline, indicating that ∼98% of cells producing Ty-1-tubulin had indeed incorporated it into the flagellum after 24h of induction.

Fig. 6A shows flagella extracts from a cell that had integrated Ty-1-tubulin along the entire length of the OF as well as the NF (Fig. 6A, +tet 24h). These flagella that incorporated Ty-1- tubulin during the initial 24h induction will be henceforth referred to as pulsed flagella. After 24- hour incubation with tetracycline, cells were thoroughly washed with medium devoid of tetracycline and then grown in the absence of tetracycline for 48 hours (5-6 cell cycles, “chase”). We then prepared samples for IFA and searched for flagella of cells that were still positive for Ty-1-tubulin, which they must have integrated during the initial 24h induction. In bi- flagellated cells which expressed tagged tubulin, we found that Ty-1-tubulin was present in the OF but not the NF, as expected (Fig. 6A, middle panel, +tet 24h, -tet 48h). This pattern is the mirror image of a short induction (Fig. 4A, Fig. S5A), which indicates that the de-induction was successful. Strikingly, Ty-1-tubulin was absent from the distal tip in 95% (n=300) of chased OF (OF in Fig. 6A, middle panel). This meant that most flagella that had been constructed during the 24h induction had integrated untagged tubulin at the distal tip during the 48h washout period.

As a last step of this experiment, we subjected the cells that were grown in the absence of tetracycline for 48h to another round of induction for 4 hours (re-induction). In the right panel of Fig. 6A a bi-flagellated cell from this timepoint is shown. It has a green OF due to the Ty-1- tubulin integration during the initial 24h induction. The NF in this cell has integrated Ty-1-tubulin in its distal segment (see case 1 scenario 2 Fig. 4A), indicating its construction had been initiated before the re-induction, evident by the proximal untagged segment. This demonstrates that the chased cells were still capable of producing and integrating Ty-1-tubulin into the NF during the 4h re-induction. However, the tagged tubulin was absent from the distal tip of the OF similar to what was observed 48h post washout. This indicates that the OF is locked in these cells, again highlighting that simultaneous integration of tubulin in both NF and OF does not occur. After de-induction Ty-1-tubulin is absent in NF and the posterior MTs of the cell body (Fig. 6A, S8B, +24 -48h, magenta arrow) as well as the tip of the OF (magenta arrows). In bi- flagellated cells that were observed after re-induction, tagged tubulin was found integrated again in the NF as well as the MTs of the posterior cell body (Fig. 6A, S8C +24 -48h +4h, green arrow).

From these experiments, we derived two important observations. First, 95% of chased OF that were labeled in the 24h induction, had integrated untagged tubulin at their distal tip, indicating that incorporation occurs regularly, most likely in all flagella. Second, these experiments (Fig. 6A) confirm our previous observations that OF of bi-flagellate cells do not incorporate tubulin at the same time as the NF (Fig. 4A), indicating that integration occurs after cell division during G1-phase.

Therefore, mono-flagellated (G1-phase) cells with a green flagellum were analysed in the re-induced conditions. Fig. 6B shows an example of a single flagellum in which multiple rounds of integration are revealed. This flagellum most likely emerged shortly before the initial 24h induction (short untagged proximal segment, labeled “before”). It was then assembled for most of its length in the presence of tetracycline (long green segment, “24h induction”). A short untagged portion then follows, which we attribute to the 48 hours in which the cells were grown in the absence of tetracycline, post washout (“de-induction”). At the very distal tip, following this untagged part, one can observe the integration of tagged tubulin again (short green segment, “Re-induction”), corresponding to the 4h re-induction. A detergent extracted cytoskeleton of a mono-flagellated cell presenting a similar pattern is shown in the right panel of Fig. S8D. Taken together, these experiments suggest that chased mono-flagellated cells had integrated tubulin at the distal tip of their flagellum again during G1-phase.

Above experiments suggest that integration of Ty-1-tubulin is observed frequently at the distal tip of OF and that this most likely occurs during the G1-phase, as it is not observed in OF of bi-flagellated cells.

### Integration at the distal tip is also found for a HaloTag-tagged component of the axoneme

Next, we wanted to corroborate these findings and exclude that they are an artifact of tubulin overexpression due to the inducible system. Hence, we employed an orthogonal system that allowed for the dynamic tracking of other axonemal components. Radial spoke protein 4/6 was N-terminally *in situ* tagged with mNeonGreen followed by a HaloTag using the PCR - tagging approach (Dean et al., Open Biol 2015), generating a culture of cells all of which expressed the tagged protein. This was validated based on presence of the mNG signal, which serves as a general reporter for RSP4/6 in these cells, regardless of time of synthesis. Although the HaloTag itself is not fluorescent, the cells can be incubated with a fluorescently labelled ligand that covalently binds the tag (Dean et al., 2016; Los et al., 2008), enabling us to perform pulse chase experiments and hence trace populations of proteins synthesized in the cell at various time points.

The cells were incubated for 1 hour with TMR ligand that can conjugate with the HaloTag. When imaged immediately after the unconjugated TMR ligand washout, both mNG signal and TMR signal localized along all flagella from their proximal to distal ends (Fig. S9A), demonstrating that the tagged protein efficiently binds the TMR-ligand. We then performed a chase experiment, when the cells were grown in a medium for two hours after the ligand washout followed by their fixation and imaging. Similarly, to the experiments with induction of Ty-1-tubulin expression (Fig. 3B), we observed that the newly synthesized RSP4/6 was incorporated into the distal portion of the new flagellum, which was manifested as a gradient of decreasing TMR intensity towards the distal tip (Fig. 7A). The OF in these bi-flagellated cells was fully labelled with TMR (Fig. 7A), supporting our earlier observations that the two flagella do not compete for building blocks (case 1 scenario 2 in Fig. 4A). When cells were fixed 4 hours after the ligand washout, bi-flagellated cells with a short NF and a decrease in the TMR intensity at the distal tip of their OF were observed (Fig. 7B), resembling results of 4-hour induction of Ty-1-tubulin expression (Fig. 4C). Moreover, we noticed that in addition to monoflagellated cells with a gradually decreasing TMR intensity along their flagellum, other cells had a high TMR intensity along the flagellum with a sharp decrease in the distal part (Fig. 7C, white arrows), mirroring the type 3 flagella in Fig. 5B.

**Fig. 7:**
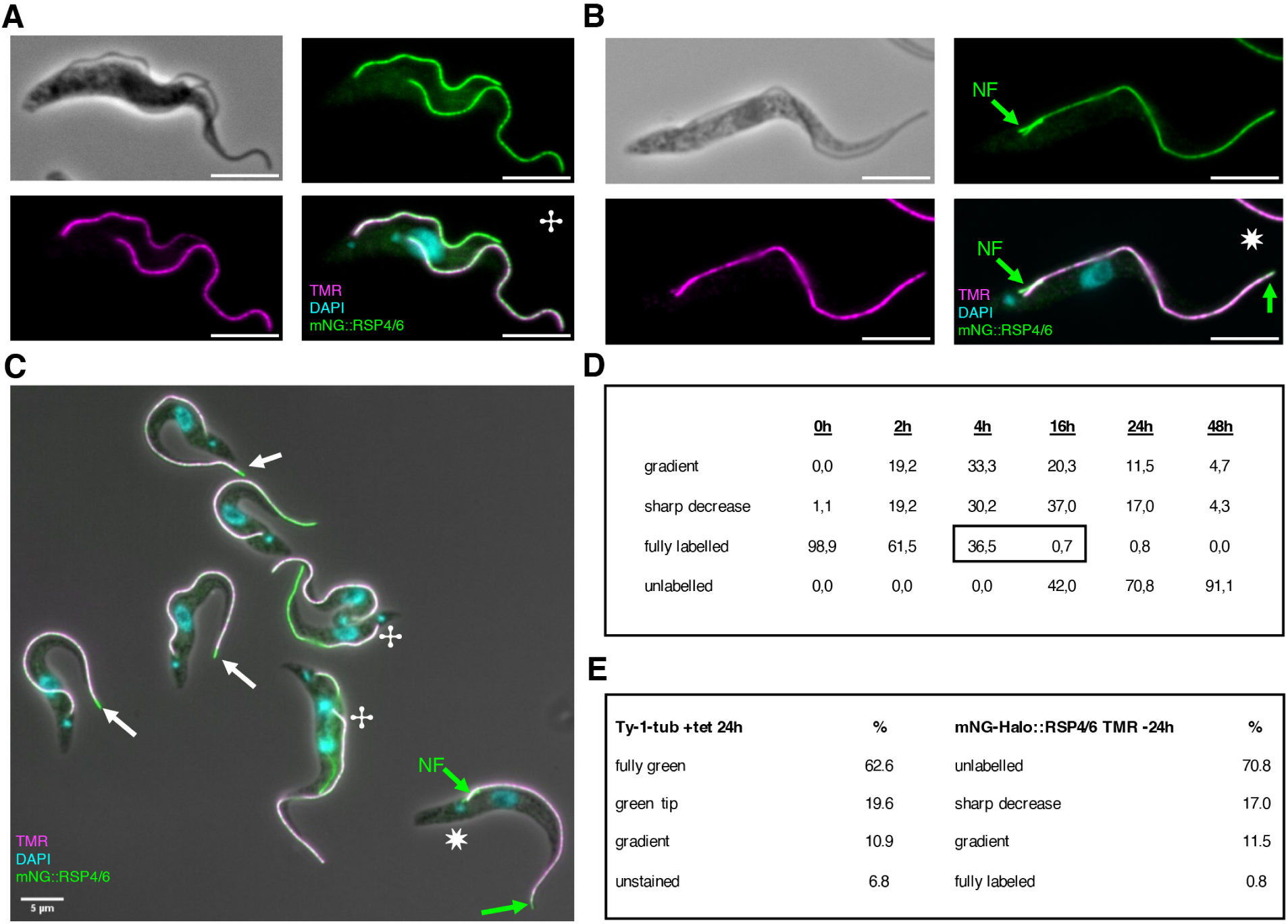
**Incorporation at the tip is found for other axonemal component visualized using a HaloTag** A: A biflagellar cell fixed **<u>2 hours</u>** after TMR labelling and ligand washout. Unlabelled RSP4/6 is incorporated in the new flagellum exclusively at its distal end. HaloTag TMR-ligand signal (magenta), mNG-HaloTag::RSP4/6 (green), DAPI (cyan). B: A biflagellar cell fixed **<u>4 hours</u>** after TMR labelling and ligand washout. Unlabelled RSP4/6 is incorporated at the distal tip of the OF and along the entire length of the NF. C: A field of fixed cells **<u>4 hours</u>** after TMR labelling and ligand washout. White arrows: monoflagellated cells with a visible sharp decrease of TMR ligand at the distal tip of the flagellum. Green arrows: integration of unlabelled RSP4/6 into short NF and the OF tip. ✢ = case 2 scenario 1 cell (similar to Fig. Fig. 4A); ✷ = case 1 scenario 1 cell (similar to Fig. 4B/C). D: The proportion of different types of flagella found after various timepoint post TMR ligand labelling + washout. Between 4 and 16h the proportion of flagella that did not incorporate unlabelled RSP4/6 drops to 0.7% (highlighted by the black rectangle). E: Comparison between Ty-1-tubulin cell line **<u>24h after the addition of tetracycline (n = 265)</u>** and mNG-Halo::RSP4/6 **<u>24h after TMR ligand labelling + washout (n = 253)</u>**. In all experiments, mono- flagellated and bi-flagellated cells were scored together. Scale bar = 5µm.

Strikingly, comparing the number of fully labeled flagella after 4h (∼0.5 cell cycles, 36.5%) and after 16h (∼1.7 cell cycles, 0.7%) shows that almost no cells retain fully labeled flagella once they had passed the threshold of completing one cell cycle (∼9h) (Fig. 7D). When the distribution of different types of flagella after 24h TMR washout was compared to the flagella type distribution after 24h induction with the Ty-1-tubulin cell line, it revealed that corresponding flagella types are present in similar numbers in the two types of experimental systems (Fig. 7E).

Hence, we concluded that the results obtained using these two orthogonal systems, inducible expression of Ty-1-tubulin and HaloTag-conjugated TMR ligand labelling of the radial spoke protein RSP4/6 expressed from endogenous locus, are consistent. In both systems the axonemal extension is restricted to the NF in bi-flagellated cells. However, after cells divide and the two flagella are no longer competing, remodeling occurs at the distal tip in all flagella, indicated by the disappearance of TMR-labeling, which is thereby a robust phenomenon.

Since flagellum length of *T. brucei* is remarkably stable, these results are only explicable with the idea of cell cycle dependent dis-assembly and subsequent re-assembly of the distal part of the axoneme.

## Discussion

A fast response rate to tetracycline (∼1h) and low level of leaky expression enabled us to elucidate incorporation of tagged alpha-tubulin into the axoneme, with enough sensitivity to determine the growth rate of the axoneme. Previously, the assembly rate of the flagellum has been measured with an extra-axonemal structure, namely the PFR (Bastin et al., 1999). At a speed of ∼2.9µm/h, the assembly rate of the axoneme measured by us using tagged tubulin is similar, albeit a little slower, than the one shown for the PFR (∼3.6µm) (Bastin et al., 1999). Similar to the PFR, we observed that the axoneme was built at a linear rate. The delayed initiation of assembly of PFR in respect to the axoneme together with the coincidental finish of their assembly is a possible explanation for the slightly higher rate at which new PFR material is added. Alternatively, this difference may be the consequence of variability between the two different cell lines in which the growth rates were measured.

In contrast to animal cells but like the vast majority of protists, trypanosomes maintain during cell division a flagellum that was constructed in one of the previous cell cycles while constructing a new one. This raises two questions: (1) how do they deal with flagella of different age? (Bertiaux and Bastin, 2020) and (2) what is the fate of each flagellum? Thanks to the data reported here, we can address both questions.

We can follow the fate of a flagellum in the models depicted in Fig. 8. First, we want to illustrate flagellum assembly by assuming the Grow-and-Lock model for a flagellum that gets locked and is then sealed from further tubulin incorporation forever (Fig. 8A). The assembly of a NF, here represented with an orange line, occupies approximately 50-60% (∼ 5h) of the cell cycle (Sherwin and Gull, 1989; Woodward and Gull, 1990). With an assembly rate of ∼2.9µm/h, this is sufficient for a flagellum to reach ∼15 µm in length (∼80% of full length). The cell then divides and the NF is assembled to full length post division during G1-phase. Based on our experiments in Fig. 5B we assume that elongation after cell division continues at a linear rate and most likely concludes soon after cytokinesis and before the assembly of a NF is initiated. (Abeywickrema et al., 2019). The flagellum then acquires CEP164C at its base, before it becomes an OF for the first time, (Atkins et al., 2021) to establish the lock. From that point on, it no longer elongats the axoneme (indicated by the red lock in Fig. 8). OF length is maintained at ∼20µm and does not change, indicated by the straight orange line. If these OF do not incorporate new building blocks for the rest of the lifespan of the cell, their proportion in a culture at any given moment is at least 50% (OF from bi-flagellated cells (∼33.3%) plus locked flagella from monoflagellated cells (16.6%)).

**Fig. 8:**
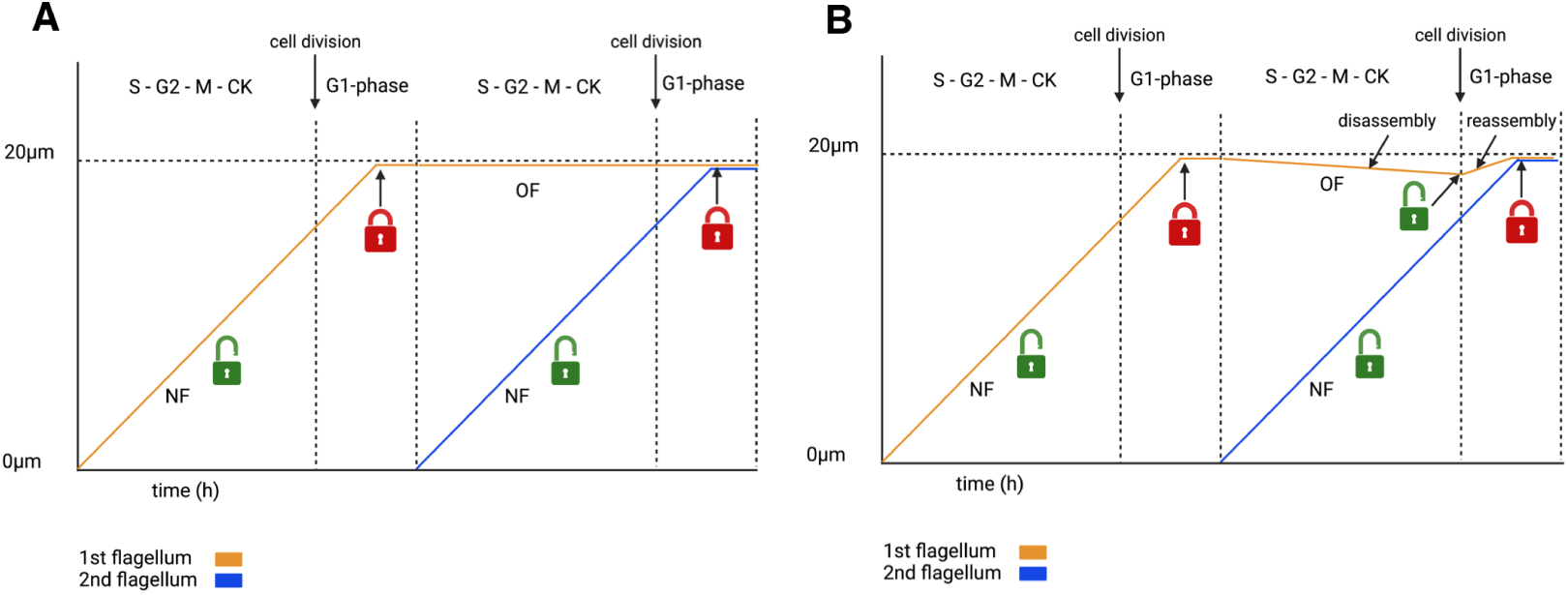
**Flagellum dynamics over multiple cell cycles** Model describing flagellum growth and maintenance in *Trypanosoma brucei* assuming the Grow-and-Lock model. A: In this graph the locking mechanism persists indefinitely. Once the maximum length is reached it remains stable and the flagellum neither shortens nor elongates. The orange line represents flagellum length, with a NF assembling at a linear rate that reaches maximum length in the following G1-phase and then gets locked. The blue line represents linear assembly of the NF inside the cell that carries the OF that is represented by the orange line. B: The flagellum assembles at a linear rate and gets locked before a NF is assembled in the same cell, represented by the blue line. While the NF assembles the OF (orange line) slightly shortens. After the cell divides the NF daughter (blue line) assembles its flagellum to full length, meanwhile the OF daughter is unlocked briefly, to allow the previously dis-assembled portion to re-assemble.

This hypothesis could be tested with the inducible expression of Ty-1-tubulin. According to the length of the cell cycle, after 9h post tetracycline addition, 50% of flagella should be devoid of Ty-1-tubulin, after 18h it should be 25%, and after 27h 12.5%, respectively. However, in reality, 98% of flagella contained Ty-1-tubulin 24 hours after the addition of tetracycline (Fig. S7). Furthermore, in the de-induction experiments, >95% of the pulsed green flagella that were chased 48h post washout had incorporated untagged tubulin again. Should flagella stay locked indefinitely, this could only be possible for the 50% of flagella that did not finish assembly during the last cell cycle of the Ty-1-tubulin labeling step. In re-induced mono-flagellated cells, two segments could be observed at the distal flagellum tip and correlated with timely distinct instances of integration (washout and re-induction, Fig. 6). Multiple cell cycles must have passed since the emergence of these cells, showing that a flagellum does not stay locked indefinitely but instead undergoes regular integration of new tubulin specifically at its distal tip. Employing an orthogonal approach with the HaloTag system for tracing RSP4/6 integration demonstrated that after less than 2 cell cycles, virtually all flagella possessed a segment containing recently synthesized RSP4/6.

To account for these observations, we propose a refined model presented in Fig. 8B: A new flagellum is assembled at a linear rate to ∼16µm and then finishes assembly post cell division. It then gets locked and another NF emerges (indicated by the blue line, Fig. 8B). While this NF elongates, the OF can no longer incorporate new building blocks, as shown here in multiple experiments. *T. brucei* most likely employs this strategy to prevent competition between the two organelles (Atkins et al., 2021; Bertiaux et al., 2018).

Although locked for growth the OF could still be shortening during elongation of its NF counterpart. Measuring the length of flagella over the course of a cell cycle indicated that OF are on average shortest in post mitotic cells (Abeywickrema et al., 2019). This leads to a conundrum: if there is such shortening in each cell cycle, it would over the course of many cell cycles result in significantly shorter flagella, something that has not been described. We propose that the previously locked OF gets unlocked (green lock, Fig. 8B), granting access for new building blocks to facilitate re-assembly of the flagellum to its full length after shortening. This could happen in a short time-window around the beginning of G1-phase, when OF and NF are not competing. This idea is supported by absence of the locking protein Cep164C in 60% of mono-flagellated cells (Atkins et al., 2021).

Taken together, our results demonstrate regular integration of new building blocks at the distal tip of the axoneme. As the length of the *T. brucei* flagellum is remarkably stable, this would require a disassembly mechanism to avoid excessively long flagella over multiple cell cycles. Although not previously reported for *T. brucei*, the flagellum shortening via disassembly of cytoskeletal structures was reported in related kinetoplastids, such as *Leishmania mexicana* (Wheeler et al., 2015), the firebug trypanosomatid *Leptomonas pyrrhocoris* (He et al., 2019), and the closely related *T. congolense* (Peacock et al., 2018). In *T. brucei*, there are certainly protein candidates located at the flagellum tip, which belong to families capable of tubulin removal. *T. brucei* kinesin 13-2 is found at the tip of both NF and OF. RNAi mediated knockdown has resulted in flagella that were constructed too long (Blaineau et al., 2007), but this result could however not be reproduced when the protein was knocked out (Chan and Ersfeld, 2010). The *T. brucei* genome encodes five putative proteins of the katanin family, which are generally known as MT severing enzymes, although this function was not yet demonstrated in T. brucei. Here they are localized to the distal tip of the flagellum (Tb 927.11.3870) or the posterior cell tip, where new tubulin is added to MTs (Casanova et al., 2009). It is conceivable that these enzymes can effectively shorten the old flagellum after tubulin integration is inhibited by the locking mechanism.

The biological relevance of this supposed turnover remains elusive: The integration of newly synthesized material is restricted to the flagellum part, that extends past the anterior cell body. This portion of the flagellum is likely more dynamic, as disassembly of the part attached to the cell body would involve remodeling of the flagellum attachment zone and might entail substantial rearrangement of parts of the cell body. The free part of the flagellum might be subject to physical forces of different magnitudes when compared to the attached part of the flagellum, which could lead to damage that requires repair by regular axonemal renewal. Moreover, the distal tip of the flagellum acts as a signaling hot spot (Bachmaier et al., 2022; Langousis and Hill, 2014; Lopez et al., 2015; Shaw et al., 2022, 2019). A cell cycle dependent restructuring of this region could allow trypanosomes to replenish components of signaling pathways that are depleted over a cell’s lifespan. Furthermore, the trypanosomes could change the composition of the tip, to adjust to different environments and abide cell differentiation.

In summary, we provide the first formal evidence for the Grow-and-Lock model in bi-flagellated cells. However, the flagellum is more dynamic than initially thought, as we also observed cell cycle dependent turnover at the distal tip of the flagellum, which occurs most likely during G1- phase. We propose a model where procyclic trypanosomes lock the OF before assembling a NF to avoid competition for building blocks. Towards the end of NF elongation, the OF slightly shortens followed by a short phase of re-assembly in G1-phase after cytokinesis. This repeats every cell cycle. The cells therefore maintain flagellum length in a manner that resembles a balance-point model but with timely distinct (cell cycle dependent) instances of disassembly and re-assembly, which are restricted to the free portion of the flagellum at its distal tip. In contrast, Chlamydomonas combines a constant dis-assembly rate with a variable assembly rate to regulate flagellum length. Here, the balance-point is reached when dis-assembly and assembly rates are equal.

Finally, we provide a robust tool to measure assembly dynamics of MTs with tubulin directly. Flagellum assembly was the main focus of this work; however, inducible expression of Ty-1- tubulin can also be used to track tubulin integration into the MTs of the cell corset and the mitotic spindle.

## Material and methods

### Cell culture

Procyclic trypanosome strain 427 were cultured at 27° in SDM-79 medium (GE Healthcare Cat No.: SH3A8023.01) pH 7.4, complemented with 10% fetal bovine serum and Hemin (7.5 mg/l). For growth curves, cells were counted every 24h with a Luminex® Guava® Muse® flow cytometer and then diluted to 2 x 10^6^ cells/ml.

To generate the mNG-HaloTag tagged cell line the SmOxP927 strain (Poon SK, Peacock L, Gibson W, Gull K, Kelly S. A modular and optimized single marker system for generating Trypanosoma brucei cell lines expressing T7 RNA polymerase and the tetracycline repressor. Open Biol. 2012 Feb;2(2):110037.) was used.

### Vectors and transfection

For in situ tagging of alpha-tubulin, one (1xTy-1) or two (2xTy-1) epitopes of the Ty-1-tag were inserted after threonine 41 inside the first 500bp of the alpha-tubulin ORF. The fragment was synthesized with XhoI and ApaI flanking sites and cloned into p2675 (reviewed in Kelly et al. 2007) by GeneCust (Boynes, France) (see Fig. S1). The resulting vectors were linearized with FspAI (4891–4898bp of p2675-alpha-TubTy1) before transfection. For inducible ectopic expression, one (1xTy-1) epitope of the Ty-1-tag was inserted after threonine 41 inside the full length alpha-tubulin ORF. (Sirajuddin et al., 2014). This fragment was synthesized and cloned between HindIII and BamHI sides of a pHD430-mNeonGreen-Ty1-IFT172 (pHD 430 derivative, Wirtz and Clayton 1995) by GeneCust (Boynes, France) (see Fig. S1). Prior to transfection the plasmid was linearized with NotI. For transfection, 4 x 10^7^ cells were harvested and then transfected with 10 µg of linearized plasmid using an Amaxa nucleofector II (Burkard et al. 2011). To obtain the α-Ty1-tubulin cell line, two rounds of transfection were carried out. 4 x 10^7^ cells were transfected with the pHD430 1xTy1-tub and then selected with phleomycin (2.5 µg/ml) for 10 days. The upcoming wells were harvested and screened for the presence of tagged tubulin by IFA using the BB2 antibody (Bastin et al., 1996). Positive wells were cloned out with serial dilutions and screened in similar fashion. Clones with expression of the tag in >90% of cells were then transfected as described above with the pHD360 (Wirtz and Clayton, 1995) to provide the tet repressor.

### Tagging

To generate the cell line expressing RSP4/6 (Tb927.11.4480) N-terminally tagged at an endogenous locus with mNG-HaloTag the PCR only tagging approach was employed (Paterou et al., 2023) using pPOTv4 containing a sequence for mNG-HaloTag and blasticidin resistence as a template and the following long primers: Tb927.11.4480 FW GTGAAGTGGAGGTGGAAAGGAGAGCCAGTTGGAGAGAAAAACCACGTACACAAGTTTTACAGAATAAATCAAGGTCAGATgtataatgcagacctgctgc Tb927.11.4480 RV

TCCTTATTCGCGCACATGAGATATGCCTTTGTCCTTGCGAACATCGCCTCAAGATCCTGTGGTTCAGGATTTACTGGCATactacccgatcctgatcc

Cells were electroporated with the PCR product using the BTX ECM 830 electroporator and 16 hours later blasticidin was added to the final concentration of 20 µg/ml. After selection of blasticidin-resistant clones genomic DNA was isolated and correct integration of the PCR product validated by PCR.

### TMR labelling and imaging

Live cells expressing mNG-HaloTag tagged RSP4/6 were incubated for 1 hour at 28°C with HaloTag TMR Ligand (Promega, G8251) 1000-fold diluted in medium. Subsequently, three washes with fresh medium were performed to remove ligand not bound to the HaloTag. Thereafter, cells were incubated in medium for a desired length of time, washed with PBS, settled on glass slides, and fixed in -20°C methanol for 20 minutes. The cells were then rehydrated in PBS for 20 minutes, mounted into 90% glycerol supplemented with 1% 1,4- antioxidant diazabicyclo[2.2.2]octane and 100 ng/ml of 4’,6-diamidino-2-phenylindole (DAPI) and imaged using the Zeiss AXIOPLAN 2 microscope with a 40X Plan-NEOFLUAR, 1.3 Oil, and 100X Plan-APOCHROMAT, 1.4 Oil objective lenses. Image acquisition was performed using a Zyla 4.2 sCMOS camera (ANDOR) and Micro-manager software. Images were processed and analysed using the Java based program ImageJ (Schindelin et al., 2012).

### Immunofluorescence assays

Procyclic cells were harvested at densities between 5 x 10^6^ – 1 x 10^7^. For whole cell analysis, cells were transferred onto poly-L-lysine coated slides (epredia, REF J2800AMNZ, Gerhard Menzel GmbH), left to air-dry for 10 – 15 minutes and then fixed with methanol at -20°C for 5 min. To extract cytoskeletons, cells were settled on poly-L-lysine slides for 10 min, then treated with 0.5% Nonidet P-40 in PEM buffer (2 mM EGTA; 1mM MgSO4; 0.1 M, pH6.9, PIPES) for 5 minutes and fixed with methanol at -20°C for 5 minutes (Robinson et al., 1991). Extraction of flagella was achieved by treating detergent extracted cytoskeletons with 1.1M NaCl twice for 5 – 10 minutes before fixation. Post fixation samples were rehydrated in PBS for 15 minutes. Primary antibodies: BB2 mouse monoclonal antibody (Bastin et al. 1996, undiluted) against Ty-1 tagged alpha-tubulin, FTZC rabbit polyclonal antibody ((Bringaud et al., 2000), 1:2000 in 0.1%BSA in PBS or BB2 hybridoma) as transition zone marker, Mab 25 mouse monoclonal IgG2a (Dacheux et al., 2012), 1:4 in 0.1% BSA in PBS or BB2 hybridoma) as an axonemal marker. Secondary antibodies: α-mouse IgG1 coupled to Alexa Fluor488 (AF488) (Jackson ImmunoResearch Laboratories, #Cat: 115-545-205), α-mouse IgG2a coupled to Cy5 (Jackson ImmunoResearch Laboratories, #Cat: 115-175-206), α-rabbit coupled to AF647 (Jackson ImmunoResearch Laboratories, #Cat: 711-605-152). DNA was stained with 4′,6-diamidino-2- phenylindole (DAPI, 10 µg/ml in PBS) for 1 minute. Glass cover slips were mounted with ProLong^TM^ Gold antifade reagent (P36930, invitrogen by Thermo Fisher Scientific). Images were acquired with a 100x objective of the Leica DMI4000 microscope (NA=1.4, photometrics PRIME 95B^TM^ back illuminated scientific CMOS camera) and analyzed with ImageJ (Schindelin et al., 2012).

### Western blot analysis

Procyclic cells were harvested at densities between 5 x 10^6^ – 1 x 10^7^. To obtain whole cell lysates cells were resuspended in 100 µl hot Laemmli buffer (Laemmli, 1970) and then boiled at 100°C for 5min. For cytoskeletal fraction the cell pellet was treated with 1% NP-40 in PEM buffer containing protease inhibitor (Roche cOmplete Mini EDTA free, #Cat: 11826153001) for 2 minutes. The pellet was harvested by centrifugation and then resuspended and boiled in Laemmli buffer for 5min at 100°C. To obtain the soluble fraction, the supernatant after whole cell lysis and removal of the cytoskeletal fraction was mixed with 2x hot Laemmli buffer and then boiled at 100°C for 5 minutes. Samples were separated by Sodium-dodecyl sulfate-polyacrylamide gel electrophoresis (SDS-PAGE) in Mini-PROTEAN TGX Precast gels from BIO-RAD (7.5% cat: #4561021 or 4 – 15% gradient gel cat: #4561083) with the BIO-RAD running system in Tris / Glycine / SDS – buffer (BIO-RAD cat: #1610772). Proteins were then transferred onto Nitrocellulose or PVDF membranes (Trans-Blot Turbo Transfer Pack BIO-RAD cat: #1704158 and #1704156) using the Trans-Blot Turbo transfer system of BIO-RAD. Membranes were blocked at room temperature in 5% milk in PBS for 1 hour. Primary antibodies diluted in 2.5% powder milk in PBS: BB2 to detect Ty-1-tubulin (mouse monoclonal, 1:10), Yl1/2 to detect tyrosinated tubulin (rat monoclonal 1:1000, (Kilmartin et al., 1971) obtained from Thermo Fisher scientific cat: # MA1-80017); TAT-1 to detect alpha-tubulin (mouse monoclonal 1:500, (Woods et al., 1989); C3B9 1:50 (Woods et al., 1989), and 6-11B- 1 (mouse monoclonal IgG2b, invitrogen Thermo Fisher Scientific cat:# 32-2700) to detect acetylated tubulin. Incubation for 1 hour at RT or 4°C overnight. Thereafter membranes were washed 3x in 0.1% PBS-Tween. Secondary antibodies were added in 2.5% milk in PBS: ECL^TM^ Anti-mouse IgG, Horseradish Peroxidase-Linked (Amersham^TM^ from Cytiva, #Cat: NA931-100 µl, 1 : 10’000), Anti-rabbit IgG, Horseradishperoxidase-Linked (Amersham^TM^ from Cytiva, #Cat: NA934-100 µl, 1 : 10’000), Anti-rat IgG, Horseradish Peroxidase-Linked (Amersham^TM^ from Cytiva, #Cat: NA935-1 ml, 1 : 10’000) and the blots incubated at room temperature for 1 hour. Membranes were then washed in 0.1% Tween in PBS 3 times for 10 minutes. The blots were then incubated with ECL Select^TM^ Western Blotting Detection Reagent for 5 minutes and revealed with an Amersham^TM^ ImageQuant 800. Protein amounts were estimated with the ImageQuantTL® software.

### Tetracycline induction and washout of the inducible Ty-1-tubulin cell line

To induce Ty-1-tubulin expression tetracycline (1µg/ml) was added to a culture flask containing cells at densities between 2 x 10^6^ – 8 x 10^6^. For time courses tetracycline was added at different time intervals to a separate flask per timepoint. Samples were then harvested at the same time by centrifugation and subject to analysis by IFA and western blot. To monitor the disappearance of Ty-1-tubulin over several cell cycles de-induction experiments were performed. A culture in logarithmic growth phase was induced with tetracycline and diluted to 2 x 10^6^ every 24h with new medium containing tetracycline. After 48h de-induction was started: the cells were washed with medium (supplemented with 10% FBS and Hemin) four times and then placed into fresh medium devoid of tetracycline. Cells were diluted every 24h to 2 x 10^6^ cells/ml and monitored by IFA at 24h, 48 and 72h post de-induction. From every time point whole cells, cytoskeletal and flagella extracts were harvested as described above and subjected to IFA.

### LC-MS/MS to asses Ty-1- tubulin acetylation

To purify flagella 3 x 10^7^ cells were harvested and washed in SDM-79 without serum for 3 times. They were then resuspended in 300 µl of 1% NP-40 in PEM buffer containing protease inhibitor (Roche cOmplete Mini EDTA free, #Cat: 11826153001) and 1 µl benzonase and then incubated for 15 min at RT. After that they were spun down and resuspended again in 1% NP-40 in PEM buffer containing protease inhibitor without benzonase. Samples were incubated at RT for 2 minutes and then spun down and washed in PEM buffer. Incubation in 1% NP-40 in PEM buffer followed by washing was repeated twice more and the samples spun down at 15000g. They were then resuspended in 1ml of 1M NaCl and incubated for 15 min. The samples were then spun down and washed in PBS and the NaCl treatment repeated one more time. After a last washing step, the samples were resuspended in water and the purity of the preparation was assessed at the microscope. The samples were then spun down and all supernatant was removed. The pellet containing purified flagella was stored at -80°C.

### Protein digestion and peptide cleanup

Proteins were denatured by adding 8 M urea in 100 mM Tris buffer (pH 7.5). Proteins were reduced using 5 mM DTT for 30 minutes at room temperature. Alkylation of the reduced disulfide bridges was performed using 20 mM iodoacetamide for 30 minutes at room temperature in the dark. Urea was diluted to below 1.5 M with 50 mM ammonium bicarbonate. Each sample was divided into three aliquots, each digested with a different enzyme. For one aliquot, sequencing-grade chymotrypsin was added at an enzyme-to-protein ratio of 1:50 (wt:wt) in 100 mM ammonium bicarbonate, and digestion was performed overnight at 37°C. For the second aliquot, thermolysin was added at an enzyme-to-protein ratio of 1:20 (wt:wt) in 0.5 mM CaCl₂, and digestion was performed for 6 hours at 65°C. For the third aliquot, sequencing-grade Asp-N was added at an enzyme-to-protein ratio of 1:10 (wt:wt) in 10 mM Tris-HCl (pH 8.0), and digestion was performed overnight at 37°C. Digestion was stopped by adding 1% formic acid. Samples were centrifuged at 16,000 × g for 10 minutes, and the supernatant containing peptides was collected. Peptide clean-up was performed using an Agilent AssayMap Bravo robot on C18 tips according to the supplier’s protocol. Peptides were resuspended in 2% acetonitrile/0.1% formic acid prior to LC-MS injection.

### LC-MS/MS analysis

The peptides were analyzed by nano-liquid chromatography-tandem mass spectrometry (LC-MS/MS) using a Proxeon EASY-nLC 1200 system (Thermo Fisher Scientific) coupled to an Orbitrap Q Exactive Plus Mass Spectrometer (Thermo Fisher Scientific). One microgram of peptides was injected onto a homemade 38 cm C18 column (1.9 μm particles, 100 Å pore size). Column equilibration and peptide loading were performed at 900 bar in buffer A (0.1% formic acid). Peptides were separated using a multi-step gradient: 3–6% buffer B (80% acetonitrile, 0.1% formic acid) in 3 minutes, 6–31% buffer B in 67 minutes, 31–62% buffer B in 25 minutes, 62–100% buffer B in 5 minutes, followed by 100% buffer B for 5 minutes and re-equilibration at 3% buffer B for 3 minutes, at a flow rate of 250 nL/min. The column temperature was maintained at 60°C. Mass spectrometry (MS) data were acquired using the Xcalibur software in data-dependent acquisition mode. Full MS scans were acquired at a resolution of 70,000, while MS/MS scans (fixed first mass at 100 m/z) were acquired at a resolution of 17,500. The AGC target and maximum injection times were set to 3 × 10⁶ and 20 ms for the survey scans, and 1 × 10⁶ and 60 ms for the MS/MS scans, respectively. The 10 most intense precursor ions were selected for fragmentation (Top 10 method) with a dynamic exclusion of 45 seconds. The isolation window was set to 1.6 m/z, and a normalized collision energy of 28 was used for higher-energy collision dissociation (HCD) fragmentation. The underfill ratio was set to 1.0%, corresponding to an intensity threshold of 1.7 × 10⁵. Precursor ions with unassigned charge states, or with charge states of 1, 7, 8, or >8, were excluded. The peptide match option was disabled.

### Data analysis

Acquired raw files were analyzed using MaxQuant software version 2.1.4.0 (Cox and Mann, 2008) with the Andromeda search engine (Cox et al., 2011; Tyanova et al., 2016). Samples were grouped by the type of experiment. The MS/MS spectra were searched against the *Trypanosoma brucei brucei* database (8,832 entries). All searches included the oxidation of methionine, protein N-terminal acetylation, and lysine acetylation as variable modifications, while cysteine carbamidomethylation was set as a fixed modification. Chymotrypsin, Asp-N, or thermolysin were selected as proteases, allowing for up to two missed cleavages. The minimum peptide length was set to five amino acids, and the peptide mass was limited to a maximum of 8,000 Da. The false discovery rate (FDR) for both peptide and protein identification was set to 0.01. The main search peptide tolerance was set to 4.5 ppm, and the MS/MS match tolerance was set to 20 ppm. The "second peptides" option was enabled to account for co-fragmentation events. Protein identification required at least one unique peptide assigned to the protein group. A false discovery rate cutoff of 1% was applied at both the peptide and protein levels. The mass spectrometry proteomics data have been deposited in the ProteomeXchange Consortium via the PRIDE partner repository under the dataset identifier PXD.

## Author contribution

Conceptualization: D.A., S.B., V.V., P.B.; Methodology: D.A., M.P., L.S., E.B., R.A., M.M., V.V. S.B. P.B.; Validation: D.A., M.P., L.S., E.B., R.A., M.M., V.V. S.B. P.B.; Formal analysis: D.A., M.P., L.S., E.B., R.A., M.M., V.V. S.B. P.B.; Investigation: D.A., M.P., L.S., E.B., R.A.; Resources: P.B., M.M., V.V.; Data curation: V.V., P.B.; Writing – original draft: D.A., V.V., P.B.; Writing – review + editing: D.A., M.M., V.V., S.B., P.B., ; Visualization: D.A., M.P., L.S., S.B., V.V., P.B.; Project administration: S.B., P.B.; Funding acquisition: V.V., P.B.

## Acknowledgment

Antibodies were kindly provided by the following sources: Melanie Bonhivers, Université de Bordeaux, provided the Mab25 antibody, Frederic Bringaud, Université de Bordeaux, provided the anti-FTZC antibody. We thank Luke Rice (University of Texas Southwestern) for suggesting to tag tubulin in the acetylation loop. We thank Parul Sharma for the helpful discussions and critical reading of the manuscript. We thank Dr. Samuel Dean, University of Warwick, for providing the pPOTv4-blast-mNG-HaloTag vector. The work carried out in the laboratory of VV was supported by the Czech Science Foundation (GA CR) project no. 23- 07370S. The work carried out in the laboratory of PB was funded by La Fondation pour la Recherche Médicale (EQU202203014654), the ANR (ANR-18-CE13-0014-01), a French Government Investissement d’Avenir programme, Laboratoire d’Excellence “Integrative Biology of Emerging Infectious Diseases” (ANR-10-LABX-62-IBEID) and a PTR grant from the Institut Pasteur (PTR-474-21).

