## Supplementary figures for "A novel approach to tagging tubulin reveals MT assembly dynamics of the axoneme in *Trypanosoma brucei*"

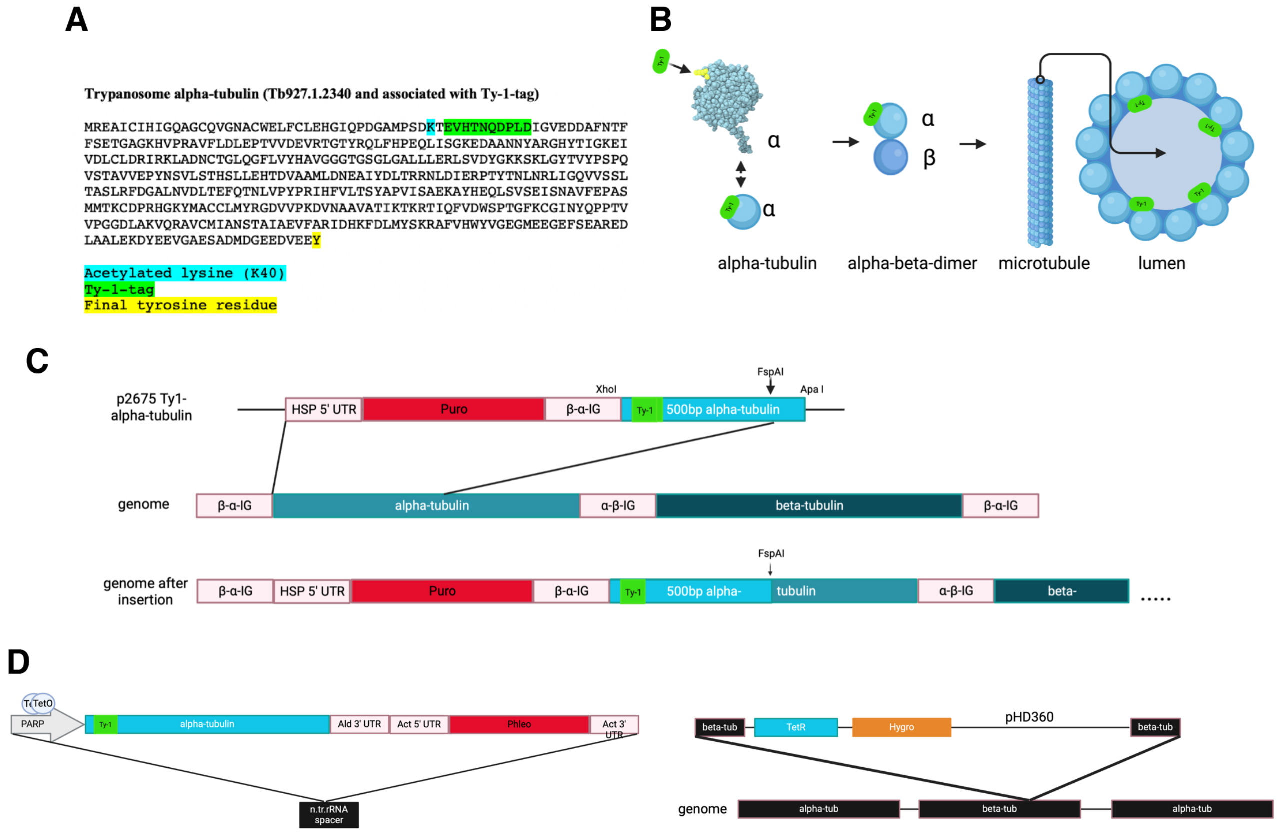


**Fig. S1: Alpha-tubulin tagging strategy**

A: Sequence of *T.brucei* alpha-tubulin. Insertion of the Ty-1-tag is highlighted in green, one amino acid after the acetylated lysine (K40, cyan). Binding epitope of the TAT-1 antibody is highlighted in red. B: Positioning of the Ty-1-tag in the acetylation loop highlighting that the tag faces the lumen of the MTs. C: In situ tagging strategy with p2675. The first 500bp of the alpha-tubulin ORF that contains the Ty-1-tag were introduced between XhoI and ApaI sites downstream of the puromycin selection marker followed by the beta-alpha-tubulin intergenic region. The tagged ORF will recombine in the genome with one of the ~40 alpha-tubulin loci under its endogenous 5'UTR (introduced by the plasmid) and its endogenous 3' UTR from the recombined tubulin locus. D: Inducible ectopic expression of Ty-1-tubulin in pHD430. The whole ORF containing the Ty-1-tag was cloned between HindIII and BamHI sites downstream of a procyclin promoter that contains two Tet operator sequences. In a first round of nucleofection 1xTy-1-tub-pHD430 was introduced. Followed by a second round of transfection with the pHD360 plasmid that introduced the Tet-repressor.

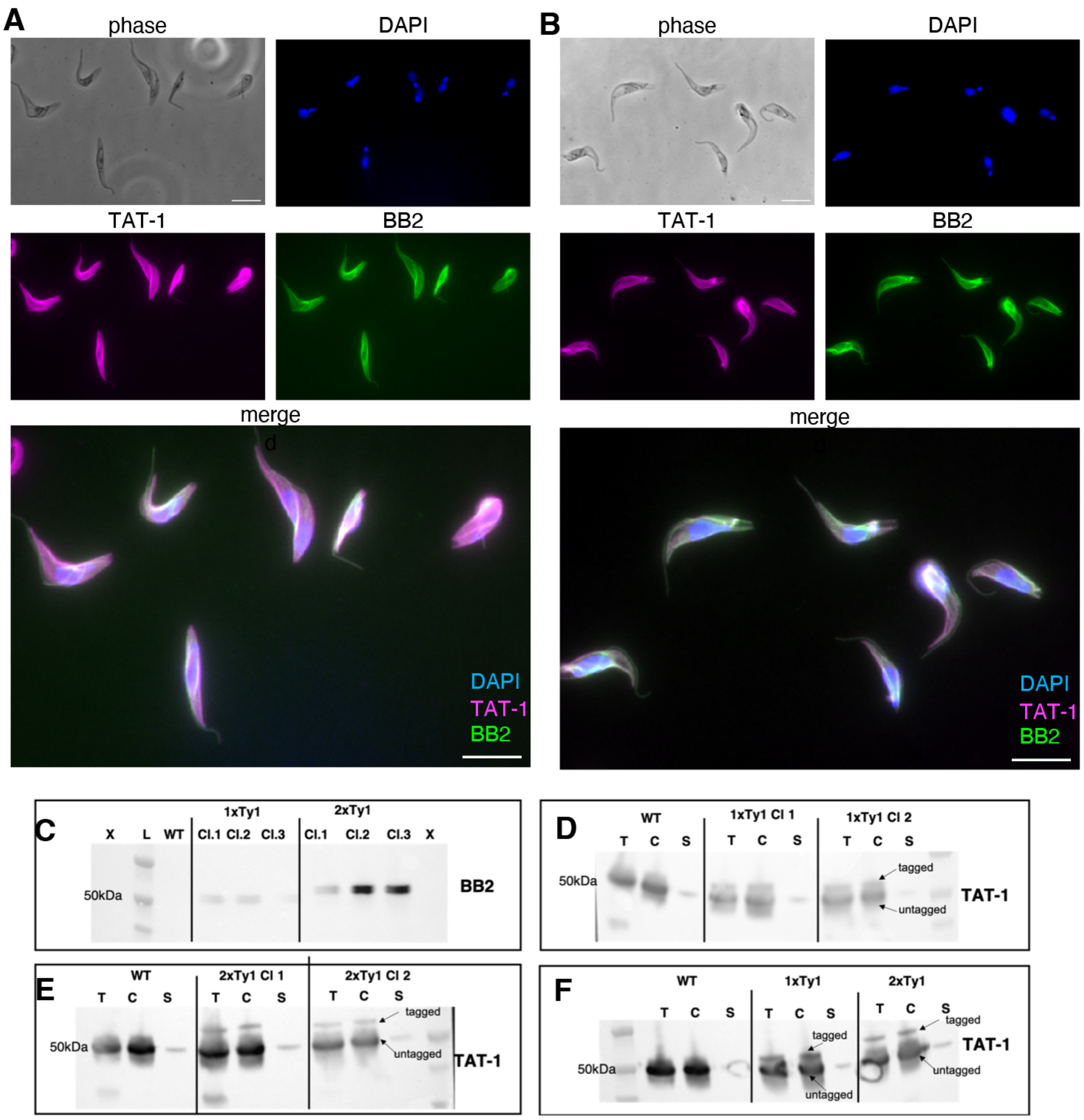


**Fig. S2: In situ tagging of alpha-tubulin**

A: Detergent extracted cytoskeletons from a cell line where alpha-tubulin was tagged with one epitope of the Ty-1-tag inside the acetylation loop (1xTy). B: Cell line that was tagged with two epitopes of the Ty-1-tag (2xTy). Samples were stained with antibodies recognising the Ty-1-tag (BB2), alpha-tubulin (TAT-1) and DAPI. C-F: Western blots with the 1xTy and 2xTy cell lines. Stained with BB2 or TAT-1 antibody. T=whole cell samples C= cytoskeleton extracts, S= soluble proteins. C: Whole cell samples from three clones of each cell line, stained with the BB2 antibody (tag only). D: T, P and S samples from two clones of the 1xTy cell line. Membrane was stained with the TAT-1 antibody. Two bands are indicated with arrows, the upper one for the tagged protein and the lower one for untagged. E: Same experiment as in D with the 2xTy cell line instead. F: T, P and S samples from one clone of the 1xTy and one of the 2xTy cell lines analysed on the same blot. Scale bar = 10$\mu$m.

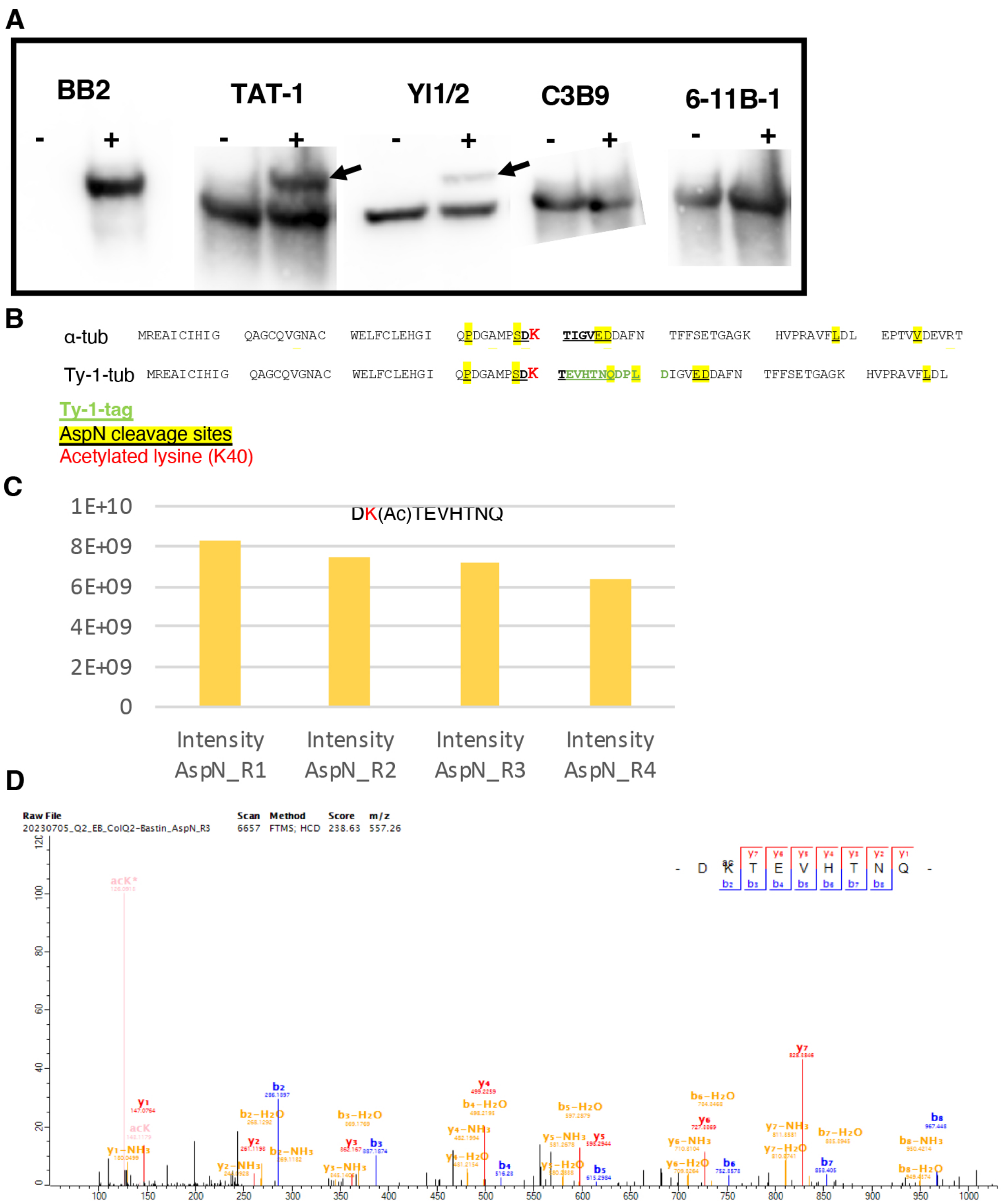


**Fig. S3: Acetylation is present on Ty-1-tubulin but cannot be detected with antibodies.**

A: Western blots with whole cell isolates from the Ty-1-tubulin cell line without (-) and 24 hours after the addition of tetracycline (+). Five individual membranes were stained with different antibodies. BB2 (Ty-1-tag), TAT-1 (alpha-tubulin), YL1/.2 (tyrosinated alpha-tubulin), C3B9 (acetylated alpha-tubulin (K40)), 6-11B-1 (acetylated alpha-tubulin (K40)). Under induced conditions a second band corresponding to Ty-1-tubulin is visible for TAT-1 and Yl1/2 but not for the two acetylation specific antibodies. B: Amino acid sequence of tagged tubulin with AspN cleavage sites marked in yellow, the acetylated lysine (K40) marked in red and the Ty-1-tag marked in green. C: MS-analysis of protein isolates from detergent extracted flagella of the Ty-1-tubulin cell line 7 days after induction. Presence of the acetylated lysine on the Ty-1-tagged peptide was found in all 4 replicates (R1-R4). D: Spectra of replicate 3 depicting different ions that were detected from the tagged peptide including the acetylated lysine (acK).

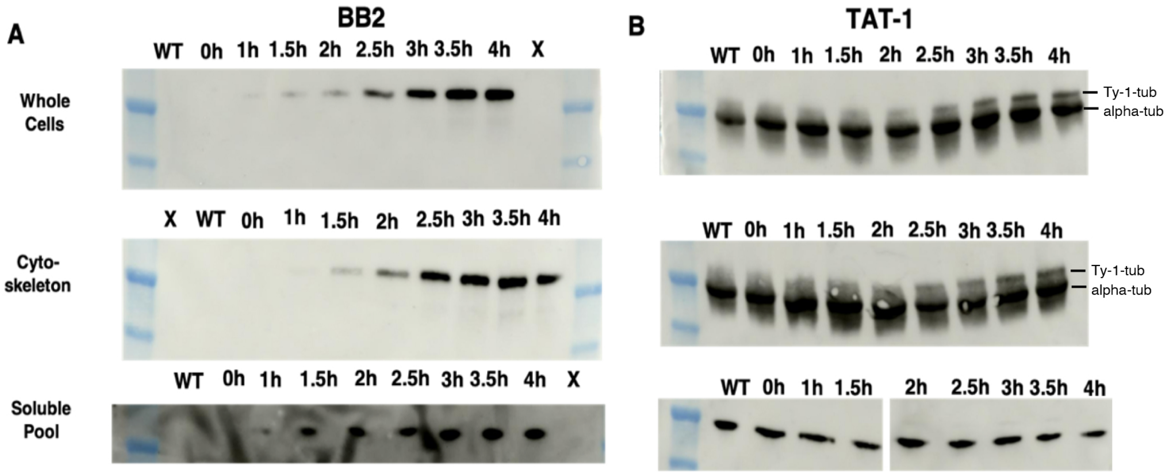


**Fig. S4: Expression of Ty-1-tubulin can be detected early after induction**

Ty-1-tubulin cell line was induced for 4 hours and cells harvested in 30-minute intervals. A: Three membranes containing whole cell isolates, cytoskeleton extracts and soluble proteins from every timepoint were stained with the BB2 antibody. B: Three separate membranes with the same samples as in S4A were stained with the TAT-1 antibody.

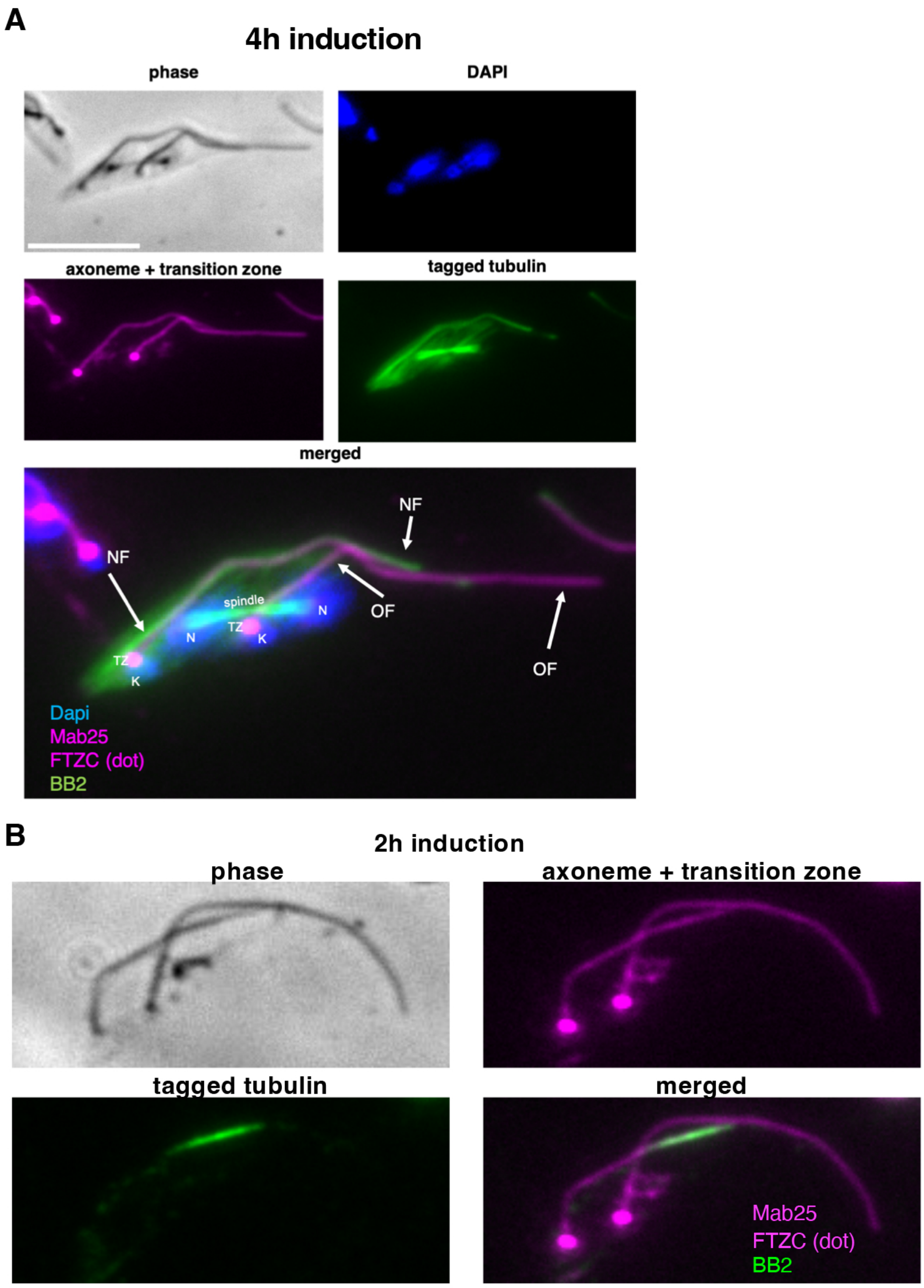


**Fig. S5: Integration of new Ty-1-tubulin.**

A: Detergent extracted cytoskeleton of a bi-flagellate cell from the inducible Ty-1-tubulin cell line after 4 hours of induction. Samples were stained with BB2 (Ty-1-tubulin, green), FTZC (transition zone, magenta dot) and Mab25 (axoneme, magenta) as well as DAPI (blue). Ty-1-tubulin has integrated in the new flagellum (NF) but not the old flagellum (OF) as well as the posterior cell body and the mitotic spindle. spindle= mitotic spindle, N = nuclei,K = kinetoplast, TZ = transition zone. B: 1M NaCl extracted flagella of a cell from the inducible Ty-1-tubulin cell line, 2 hours after induction. Samples stained with BB2, Mab25 and FTZC. Ty-1-tubulin is incorporated at the distal tip of the NF but not the OF. Scale bar = 10$\mu$m.

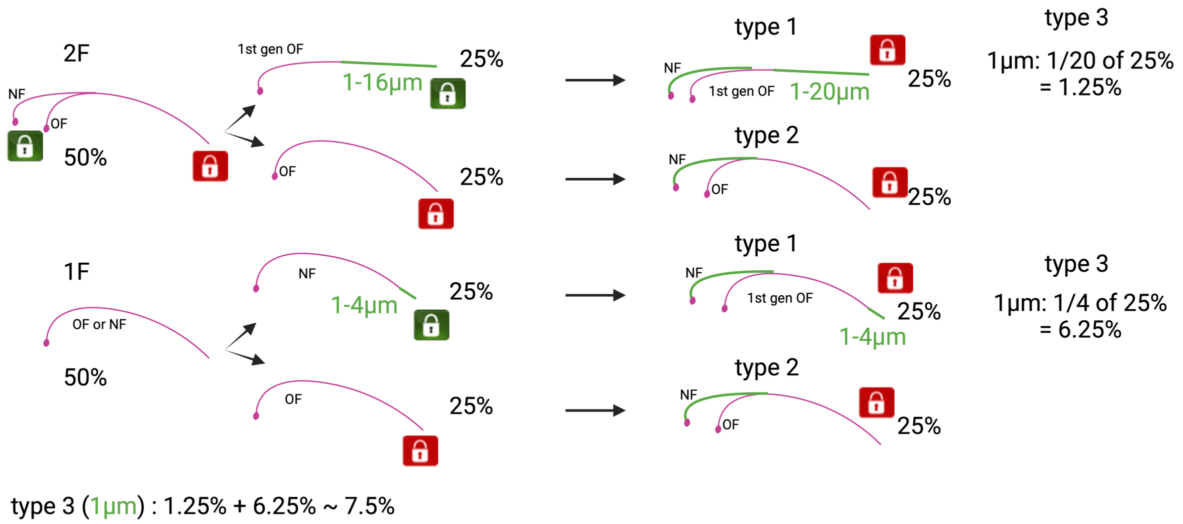


**Fig. S6: Predicted Ty-1-tubulin incorporation assuming the Grow-and-Lock model.**
One cell cycle of an average culture at densities ~2 - 8x$10^6$ cells/ml consists of ~50% mono-flagellated and ~50% bi-flagellated cells. Assuming the Grow-and-Lock-model only NF of bi-flagellated cells (2F) should integrate Ty-1-tubulin (green) at the distal tip after induction. They will integrate Ty-1-tubulin in a segment of 1 – 20$\mu$m. In the next cell cycle when these flagella have become an OF and a small subset (1.25%) will have incorporated Ty-1-tubulin in short segment of ~1$\mu$m, since assembly follows a linear rate. The OF does not integrate new tubulin as it is locked (25%). Half of the mono-flagellated cells (1F) correspond to previously OF post-division and do not incorporate new tubulin either. The other half corresponds to NF in the previous division and have completed assembly of the last ~4$\mu$m post division. Approximately ¼ of these flagella (total: ~6.25%) will incorporate Ty-1-tubulin in a segment of 1$\mu$m, assuming assembly post-division follows a linear rate. According to the Grow-and-Lock-model as well as linear assembly rate, ~50% of OF should not incorporate Ty-1-tubulin and ~7.5% should have incorporated Ty-1-tubulin in a segment of ~1$\mu$m at the distal tip.

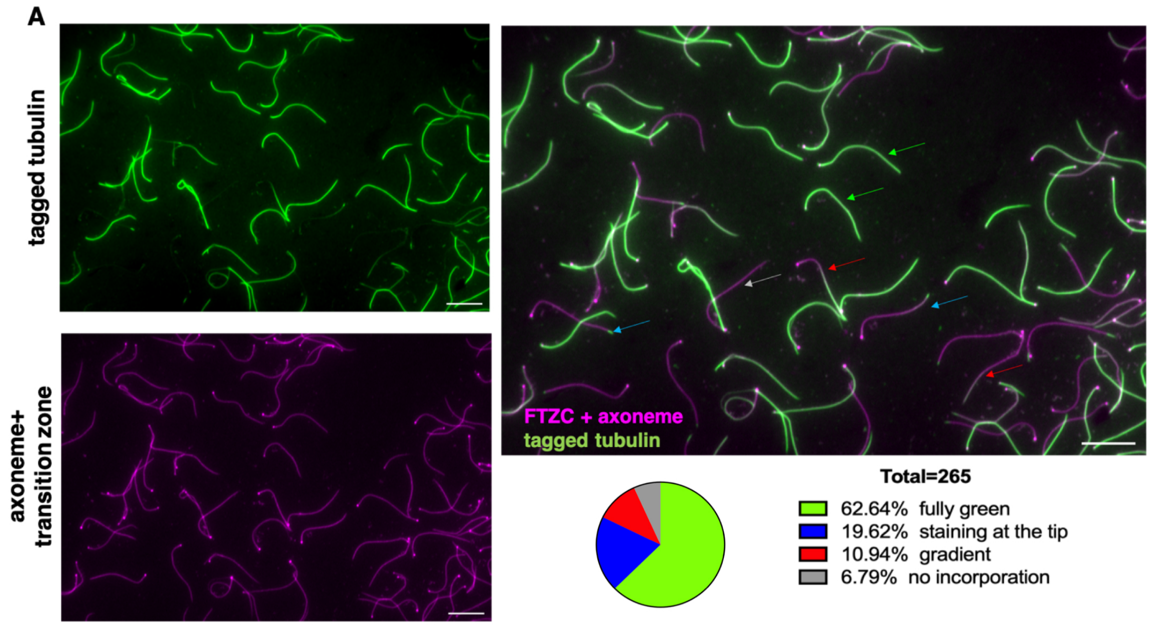


**Fig. S7: >95% of flagella incorporate Ty-1-tubulin after 24h**

A: 1M NaCl extracted flagella from the Ty-1-tubulin cell line 24h after the addition of tetracycline that were stained with BB2 (Ty-1-tag), Mab25 (axoneme) and FTZC (transition zone). Arrows indicate four types of flagella: green= flagella that incorporated Ty-1-tubulin along the entire length, red= flagella with a gradient, blue= flagella with Ty-1-tubulin at the distal tip, grey= flagella that did not incorporate Ty-1-tubulin. In whole cells from the same experiment 5% of cells (total n = 265) did not incorporate any Ty-1-tubulin in the cell body. The pie chart on the right depicts the relative proportion of different types of flagella found after 24h of induction. Scale bar = 10$\mu$m.

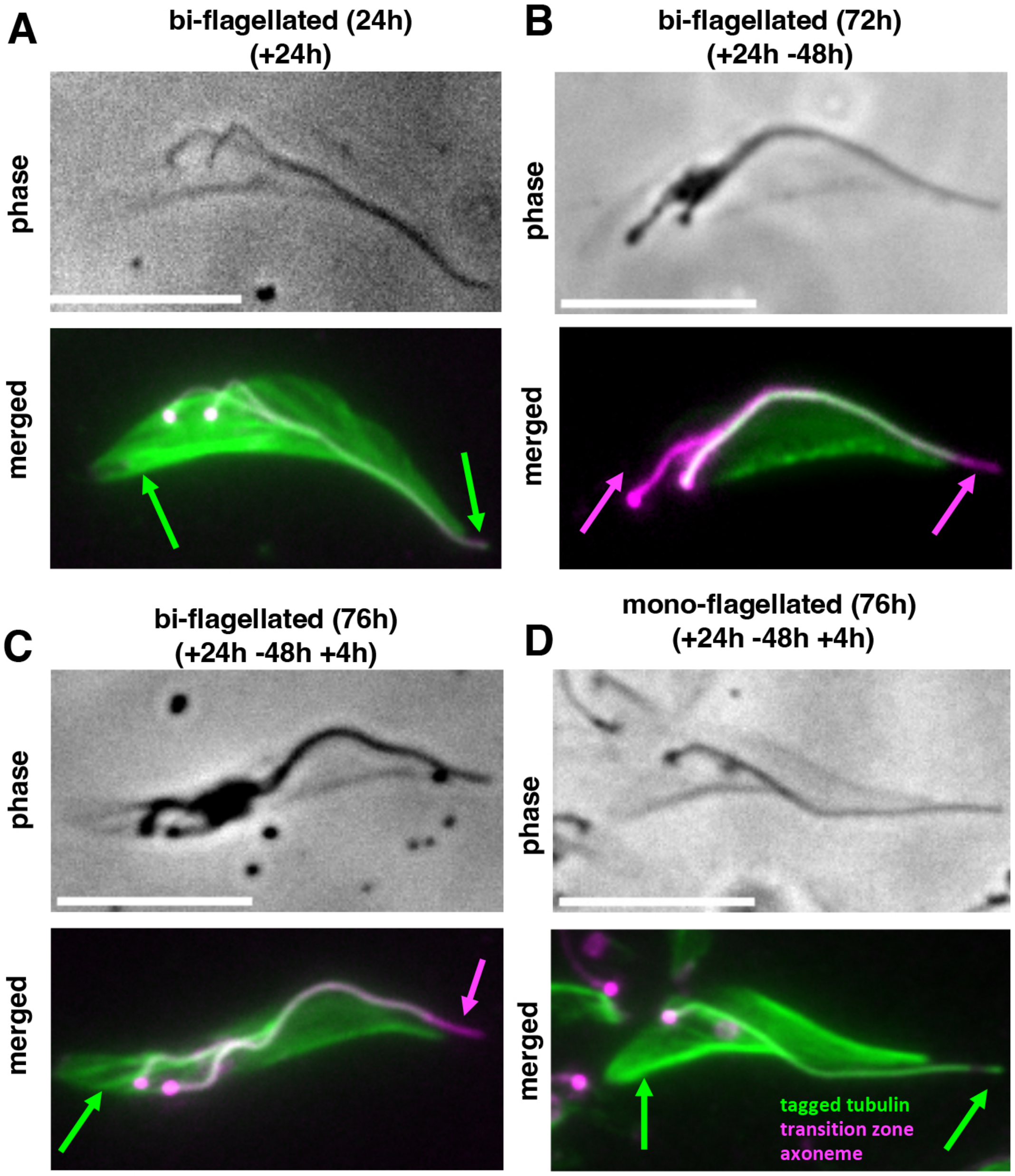


**Fig. S8: Detergent extracted cytoskeletons of de-induction experiments**

Detergent extracted cytoskeletons from the same experiment as in Fig. 6. Green arrows highlight presence of Ty-1-tubulin in posterior cell body, NF, and flagellum tip. Magenta arrows indicate Ty-1-tubulin absence, respectively. A: After 24h of induction, Ty-1-tubulin is present in NF and OF (from base to tip). B: 48h post de-induction Ty-1-tubulin is absent from the NF but present in the OF, except the OF distal tip. C: 4h after re-induction Ty-1-tubulin is present in the NF and the OF, but absent from the OF distal tip. D: Detergent extracted cytoskeleton of a monoflagellated cell. Scale bar = 10$\mu$m.

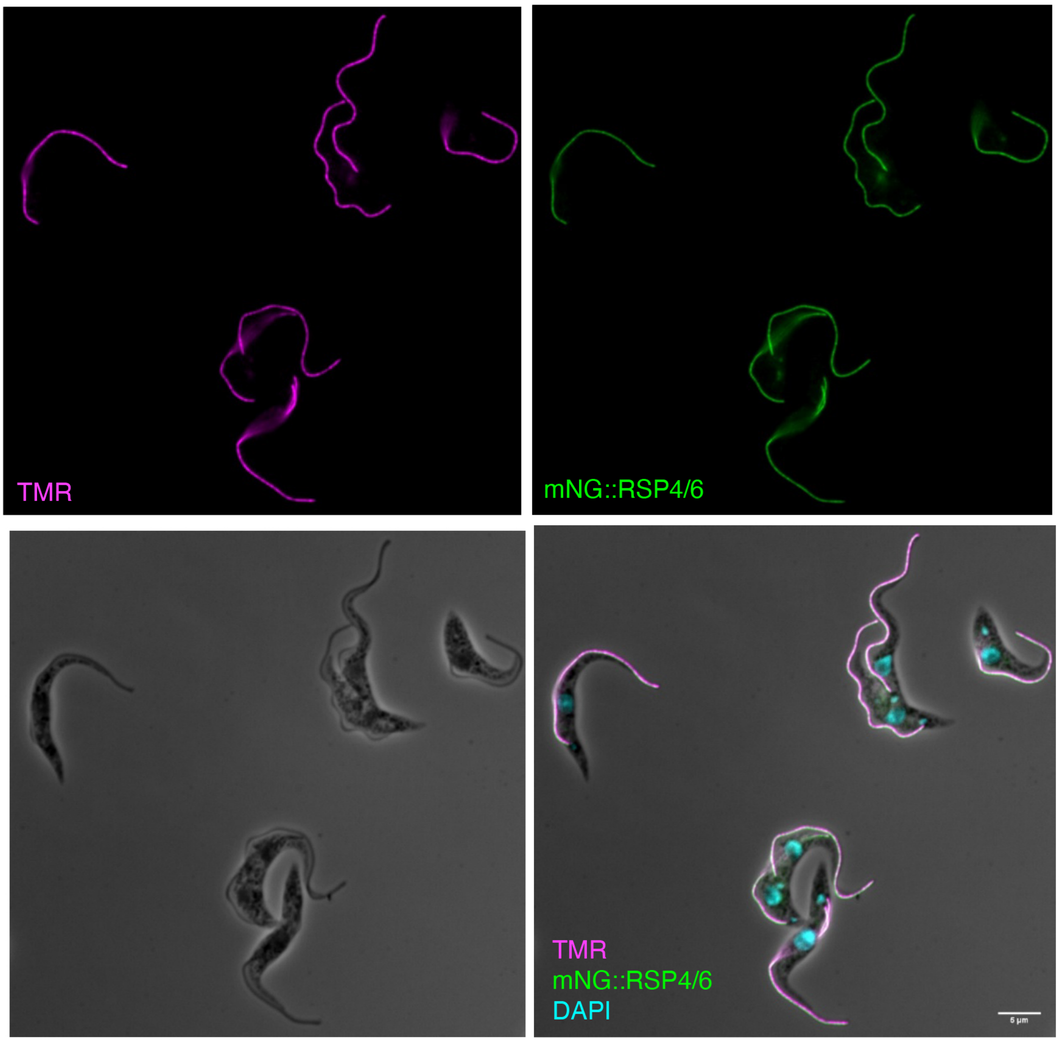


**Fig. S9: mNG-HaloTag::RSP4/6 localizes to the flagellum and is efficiently labelled with TMR-HaloTag ligand.**

mNG-HaloTag::RSP4/6 cells were incubated with TMR-ligand for 1 hour. The ligand was then washed out and the cells immediately fixed. TMR signal is shown in magenta, mNG in green and DAPI in cyan. TMR and mNG signal colocalize along the entire length of both flagella. Scale bar = 5$\mu$m.
